# Influenza virus recruits “PKC⍺-MEK1-ERK2” complex to regulate nuclear export of viral ribonucleoproteins and promote virus replication

**DOI:** 10.1101/2025.03.06.641832

**Authors:** Indrani Das Jana, Soumik Dey, Manoj Si, Arunava Roy, Arindam Mondal

## Abstract

Host kinases had long been implicated in regulating various steps of influenza virus replication cycle. Previous studies showed that the protein kinase C⍺ (PKC⍺) activated mitogen activated protein kinase (MEK)/ extracellular signal-regulated kinase (ERK) cascade promotes nuclear export of viral ribonucleoprotein complexes (RNPs), by triggering phosphorylation of viral nucleoprotein (NP), the major constituent of RNPs. But, the molecular mechanism by which PKC⍺ coordinates with specific members of the MAPK pathway to regulate NP phosphorylation remained obscure. Additionally, the molecular interactions of these kinases with NP and the spatiotemporal dynamics of such interactions have never been investigated. Here we unravel the existence of a tripartite PKC⍺-MEK1-ERK2 complex that associates with influenza virus NP and mediates its phosphorylation during the course of infection. Using an analogue sensitive kinase, we show that ERK2 can directly phosphorylate NP at specific serine-threonine residues, which promote vRNP nuclear export and are indispensable for virus propagation. PKC⍺ not only activates the MAP kinase, but acts as a scaffold for mediating stable interactions between MEK1, ERK2 and NP, thereby forming an NP associated multi-kinase complex that facilitate its phosphorylation. This multiprotein complex localizes in the nucleus early during infection but into cytoplasm at later stages and overexpression of a dominant negative PKC⍺ blocks this complex formation, NP phosphorylation, vRNP export and progeny virus production. These data not only unravel a complex network of virus-host interactions supporting different influenza A and B virus replication, but also shade light upon an unexplored avenue of PKC⍺ mediated regulation MAPK pathway.

**Significance:** The substrate specificity of kinases is the key for precise regulation of various signaling cascades and can be governed by direct kinase-substrate interaction or indirect interaction mediated through scaffolding or anchoring proteins. This study shows how influenza virus exploit a well-known cellular kinase, PKCα, as a scaffold protein to activate and recruit other cellular kinases, MEK1 and ERK2, upon viral replication machinery, RNPs. This RNP associated multi-kinase complex facilitates ERK2 mediated phosphorylation of viral NP protein, which regulates RNP nuclear export and ultimately the production of new virion particles. This work elucidates an unconventional mechanism of kinase-substrate interaction which is critical for influenza virus replication and hence paves the way towards the development of novel host directed anti-influenza therapy.

## Introduction

Influenza viruses cause respiratory illness leading to seasonal epidemics and sporadic pandemics with variable morbidity and mortality rates (1). Rapid evolution through antigenic drifts and antigenic shifts results in the emergence of newer influenza virus strains and subtypes rendering available drugs and vaccines ineffective (2, 3). Hence there is a pressing need of characterizing conserved virus-host interactions that could be targeted to develop newer host directed therapeutics.

Ribonucleoprotein particles (RNPs) of influenza viruses execute viral gene transcription and genome replication within the host cell nucleus. Eight negative sense viral genomic RNA segments, individually gets enwrapped with nucleoprotein (NP) oligomers and associates with viral RNA polymerase (RdRp) constituted of PB1, PB2 and PA subunits, together forming the viral RNPs (vRNPs) (4–7). Upon entry into the host cells, incoming vRNPs are actively transported to the nucleus within an hour post infection (hpi) (8–11). In nucleus, RNPs perform primary transcription early during infection (12). At later stages in conjunction with newly synthesized viral NP and RdRp proteins, RNPs execute viral genome replication and assemble progeny vRNPs. Newly assembled vRNPs are actively exported out of the nucleus within 5-6 hpi and trafficked to the plasma membrane to assemble into new virion particles (9, 13, 14). Viral matrix protein (M1) and nuclear export protein(NEP) together recruit host Chromosome region maintenance protein (CRM-1) to mediate nuclear export of newly synthesized vRNPs (15–17). The nuclear import-export of influenza vRNPs are tightly regulated through dynamic phosphorylation of NP at multiple serine, threonine and tyrosine residues and blocking these phosphorylation attenuates virus replication in cell culture and in vivo (18–22). Although the role of NP phosphorylation in vRNP’s nuclear-cytoplasmic trafficking has been extensively studied, the host kinases responsible for NP phosphorylation and their spatiotemporal regulation are sparsely characterized (23, 24).

The mitogen activated protein kinase (MAPK), Raf/ MEK/ ERK signalling has long been implicated to modulate influenza virus life cycle, specifically by promoting vRNP nuclear export (25–27). Previous studies showed that membrane accumulation of viral HA protein, during early and late phases of infection, leads to a biphasic activation of MAPK pathway ensuring a timely export of vRNPs during the late phase of virus life cycle (28). Supporting this notion, specific inhibition of MEK or ERK blocks vRNP nuclear export and severely restricts influenza A and B virus infection in cells and in vivo (25–27, 29–31). Recently, ERK activated p90 ribosomal S6 kinase 1 (RSK1) has been shown to directly phosphorylate NP at S269 and S392 residues to promote vRNPs nuclear export (22). The Rsk1 mediated NP phosphorylation establishes the vRNP-M1-NEP-CRM1 complex in the nucleus and mediate the export of newly produced vRNPs to the cytoplasm. Interestingly the NP S269A/S392A mutant lacking Rsk1 phospho-sites, replicated to a significant extent (10^6^ log_10_ by 32 hpi), only one log_10_ lesser compared to WT virus (22). This data indicates the presence of additional phospho-sites regulated by other host kinases that compensated for this dearth of RSK1 mediated phosphorylation in promoting vRNP nuclear export.

In addition to MAPKs, host Protein Kinase C (PKC) isoforms are also known to modulate influenza virus infection. Binding of viral HA to the cell-surface receptor activates the PKC pathway (32). PKCδ has been shown to stably associate with both NP and RNP complexes by directly interacting with viral PB2 protein thereby regulates its oligomerization and assembly of the progeny vRNPs during genomic RNA replication (33). The PKCδ can also phosphorylate RdRp subunits, which has been predicted to regulate viral transcription (34). In contrast, PKCα activates the MEK/ ERK pathway to facilitate nuclear export of newly assembled vRNPs (22, 28). Pan PKC activators and inhibitors promote exclusive cytoplasmic or nuclear localization of NP/ RNPs respectively (13). These data indicate a critical role of PKCα in moderating vRNP export, although the underlying molecular mechanism remains elusive. Whether PKCα acts just as a passive activator of the MAPK cascade, or plays a direct role in NP phosphorylation is yet to be elucidated. Furthermore, molecular interaction between these host kinases (PKCα, MAPKs) and influenza NP/ RNP complex has never been investigated.

In this study, we unravel a detailed molecular mechanism by which PKCα activates the MAPK pathway through constitution of a multi-kinase complex along with MEK1 and ERK2, which stably associate with viral NP protein and phosphorylates it to regulate vRNP nuclear export. We, for the first time show that ERK2 can directly phosphorylate influenza A H1N1 and influenza B virus NP proteins in vitro and using an analogue sensitive ERK2, we confirm this observation in the context of A/H1N1/WSN/1933 infection. This ERK2 mediated NP phosphorylation promotes vRNP nuclear export and recombinant viruses lacking these ERK2 phospho-sites are severely attenuated. Our data show that the active PKCα scaffolds a stable interaction between MEK1, ERK2 and viral NP/ RNPs, thereby resulting in the formation of the multi kinase-RNP complex within the nucleus, early during infection. The same complex gets exported out to the cytoplasm at later stages thereby dynamically regulate NP/ RNP trafficking across the nuclear membrane. Interference with the PKCα activity blocks vRNP export and attenuates virus replication, but mutant NP viruses lacking the ERK2 phospho-sites are resistant to this attenuation. Together our results elucidate a completely new unexplored avenue of host PKCα-MAPK pathway which is hijacked by influenza viruses to regulate the timely progression of infectious cycle.

## Results

### 1. PKCα activated ERK2 phosphorylates influenza virus NP

PKCα can activate the MAPK cascade by directly phosphorylating Raf-1 which in turn phosphorylates and activates its downstream kinases (Fig 1A) (35). To investigate the role of PKCα in modulating MAPK mediated NP phosphorylation, we overexpressed influenza A/H1N1/WSN NP and host ERK2 in HEK293T cells and stimulated the cells using a pan PKC activator, phorbol 12-myristate 13-acetate (PMA). We showed earlier that PMA stimulation leads to NP hyperphosphorylation, leading to the appearance of a slower migrating band in SDS-PAGE (36). Interestingly, overexpression of ERK2 elevated PMA triggered NP hyperphosphorylation by 2.5 fold, thereby suggesting a potential synergism between PKC and MAPK activities in NP phosphorylation (Fig 1B). To further validate this, we useda constitutively active PKCα catalytic domain (PKCα-CAT) (37) that can trigger robust phosphorylation and hence activation of the ERK2 in cells (Fig S1, A). Co-expression of PKCα-CAT, resulted in moderate hyperphosphorylation of NP, which gets significantly boosted in presence of ERK2, in a dose dependent manner (Fig 1B, S1 B). Considering that PKCα served as one of the upstream activators of the Raf/ MEK/ ERK cascade (Fig 1A), our data indicates that either ERK2 or one of its downstream effector kinases is phosphorylating NP. To test whether ERK2 can directly phosphorylate NP, we performed in vitro kinase assay where FLAG-tagged ERK2 was overexpressed and affinity purified from PMA stimulated HEK293T cells and was subsequently used to phosphorylate recombinant purified NP. We also purified MEK1 and MEK2 from PMA stimulated cells and tested their ability to phosphorylate NP in vitro. As shown (Fig 1C), active ERK2directly phosphorylated A/H1N1/WSN NP while MEK1 or MEK2 failed to do so. ERK2 also phosphorylates the influenza A/H1N1/PR8 and the influenza B/ Brisbane/ 60/ 2008 NP proteins in vitro to the extent comparable to the A/H1N1/WSN NP (Fig 1D). These data suggest a direct kinase-substrate relationship between the host ERK2 and viral NP protein, which is largely conserved across influenza A and B viruses.

**Fig 1.**
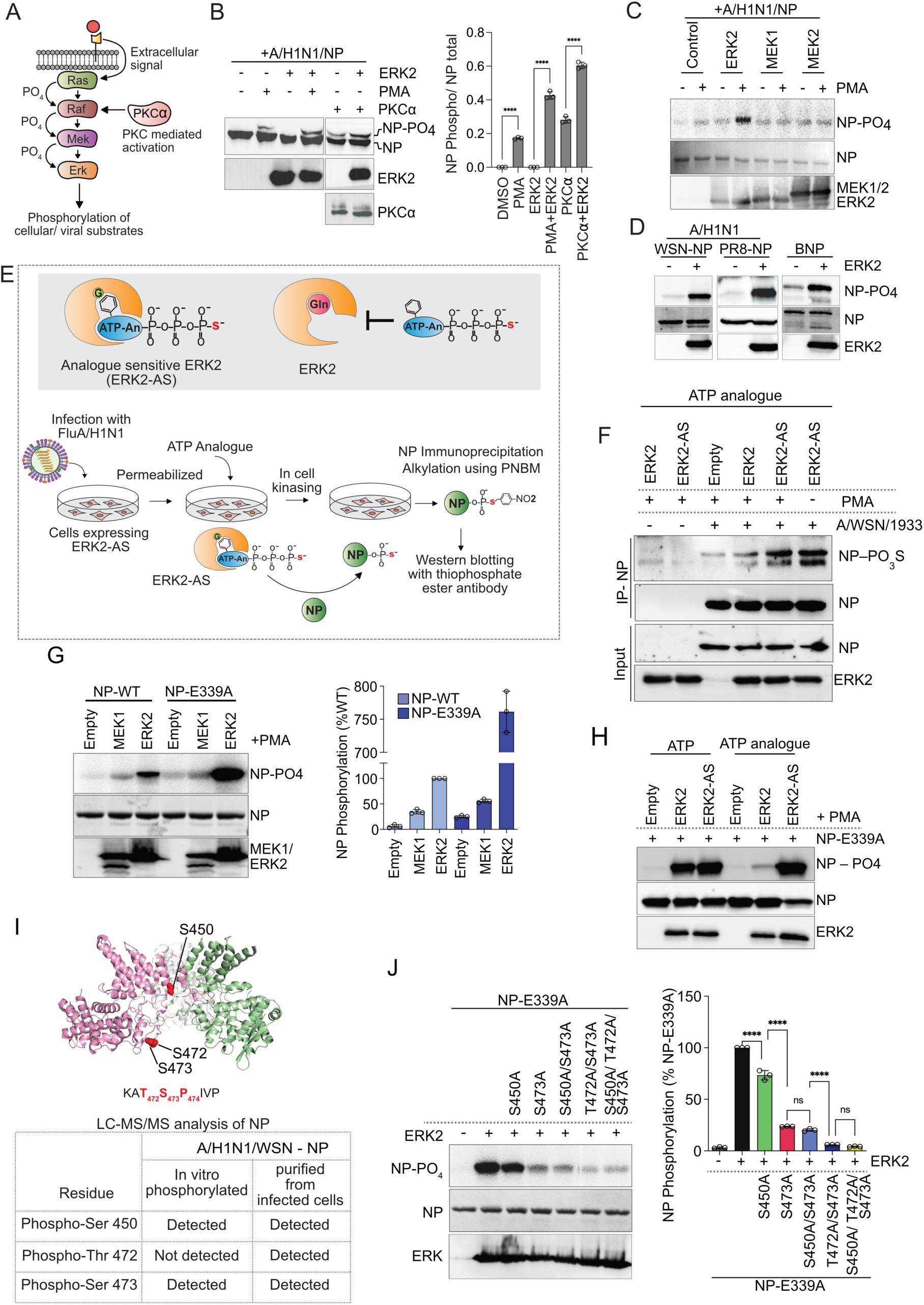
ERK2 phosphorylates influenza A NP. (A) Schematic representation of PKCα modulated ERK2-MAPK pathway. (B) HEK 293T cells overexpressing NP-V5, or co-expressing NP along with ERK2-FLAG or PKCα-CAT-HA or both were treated with PMA or left untreated. NP, ERK2 and PKCα was detected by western blotting using specific antibodies. Band intensities from three independent experiments are plotted using image J software. (C) ERK2-FLAG, MEK1-FLAG and MEK2-FLAG was purified from overexpressing HEK293T cells stimulated with PMA and used to phosphorylate recombinant purified NP in vitro. (D) In vitro kinasing of recombinant PR8 and FluB NP using ERK2-FLAG purified from PMA stimulated cells (E) Schematic representation of the mode of action of ERK2-AS and experimental design for the detection of thio phosphorylated NP. (F) Result of the experiment presented in (E). (G) WT NP or oligomerization defective NP-E339A were invitro phosphorylated using MEK1-FLAG or ERK2-FLAG. Band intensities from three independent experiments are plotted using Image J software. (H) WT NP or NP-E339A was phosphorylated invitro with ERK2 or ERK2-AS in presence of ATP or thio-ATP analogue. (I) Structure of trimeric NP (PDB ID-2iqh) showing phosphorylation sites as identified through LC-MS/MS analysis The ERK2 consensus motif present in NP primary sequence is highlighted. (J) NP-E339A harboring phosphonull alanine mutations were subjected to in vitro kinasing by ERK2. Extent of NP phosphorylation were measured from three independent experiments and plotted using image J. Two way Anova was used to measure the statistical significance between the individual sets with P value (ns > 0.05; *P ≤ 0.05; **P ≤ 0.01; ***P ≤ 0.001)*P < 0.001 denotes statistically significant.

To validate the in vitro kinase assay result in the context of virus life cycle, we adopted a chemical-genetic approach and employed the analogue sensitive ERK2 (ERK2-AS) to phosphorylate viral NP within the cell during the course of infection. The gatekeeper residue (glutamine, Q105) situated within the substrate binding pocket of the human ERK2 was mutated to glycine (G) (38) to make it compatible of utilizing the thio-ATP analogue (Fig 1E, upper panel). The HEK293T cells either overexpressing ERK2 or ERK2-AS, were infected with influenza A/WSN/1933 virus (MO1 0.01). At 12 hours post infection (hpi), the cells were stimulated by PMA, or left untreated, followed by in cell kinasing in presence of thio-ATP analogues. ERK2-AS phosphorylated NP should harbor thio-phosphate moiety conjugated at the phosphorylation sites, which upon alkylation using p-nitrobenzyle mesylate (PNBM) could be detected using thiophosphate ester specific antibody (Fig 1E lower panel). As shown, ERK2-AS catalysed thio-phosphate conjugation to the viral NP protein with high efficiency in cells infected with influenza A virus and stimulated with PMA (Fig 1F lane 5). Additionally, virus infection alone (without PMA treatment) also triggered ERK2-AS mediated NP phosphorylation to the extent comparable to the PMA treated set (Fig 1F, lane 6), thus indicating robust activation of the MAPK pathway, specifically ERK2, during influenza virus infection. Together, our data unambiguously established that influenza virus NP is a substrate for ERK2 mediated phosphorylation and suggested a role of PKCα in promoting this phosphorylation.

### 2. ERK2 phosphorylates specific serine threonine residues in NP

Next, we intend to identify the ERK2 phosphorylation sites within NP. Recombinant purified NP forms homo-oligomers (trimer/ tetramer) (39), thereby reducing the accessibility of a large number of amino acid residues, situated along its homotypic interaction interface, towards the kinases (33). Hence, to nullify any effect of NP oligomerization towards ERK2 mediated phosphorylation, an obligatory monomeric NP mutant, NP-E339A, was used as a substrate for in vitro kinasing (39). The monomeric NP-E339A appeared to be a superior substrate for ERK2, showing 7.6 fold increase in phosphorylation compared to the wildtype NP (Fig 1G), which resembled our previous observation for PKCδ mediated NP phosphorylation (33). Furthermore, the analogue sensitive ERK2-AS also showed high kinasing efficiency towards NP-E339A (Fig 1H), thereby confirming that it was ERK2, and not any other co-purified kinase,phosphorylated monomeric NP in vitro. Subsequently, this in vitro phosphorylated NP-E339A was subjected to liquid chromatography coupled with tandem mass spectrometry (LC-MS/MS) leading to the identification of serine 450 and serine 473 as ERK2 phosphorylation sites (Fig 1I, S2, Table S1-3). The S473 residue is situated within a putative ERK2 target motif (Pro-X-S/T-Pro or S/T-Pro) while the S450 residue doesn’t exhibit any such motif (Fig 1I). Viral NP purified from the influenza A/WSN/1933 infected cells, when subjected to LC-MS/MS analysis, showed high abundance of phosphopeptides harbouring phosphorylation at S450, T472 and S473 amino acid residues (Fig 1I, S3, Table S4-6), as also reported previously (36, 40). The adjacent position of T472 and S473 along with the preceding proline residue constitute the putative ERK2 phosphorylation site thus indicating that ERK2 may phosphorylate S450, T472 and S473 residues in infected cells. Interestingly, the NP T473 residue is least conserved with occurrence only in few specific strains of influenza A/H1N1, while the S450 and T472 remains largely conserved across different influenza A subtypes and influenza B virus lineages (Fig S4, Table S7).

To this endwe mutatedS450, T472 and S473 residues to phospho-null alanine, either individually or in combination, in the monomeric NP-E339A. Propensity of these mutant NP proteins were assessed towards ERK2 mediated phosphorylation in vitro. The phospho-null mutant NPs showed a varied degree of reduction in the ERK2 mediated phosphorylation (Fig 1J), however, no sensitivity was observed towards PKCδ (Fig S5), a kinase that was previously reported to phosphorylate NP at serine 165 and serine 407 residues (33). Individual mutation at S450A showed a moderate 27% decrease while S473A mutation showed drastic 76% reduction in phosphorylation compared to the control NP-E339A. Interestingly, double mutant S450A/ S473A showed reduction similar to the S473A mutant (80% reduction), hence indicating that phosphorylation at S473 is essential for subsequent phosphorylation at S450. The double mutant harboring T472A/ S473A showed a drastic 94% reduction, which is comparable to the reduction observed for the triple mutant S450A/ T472A/ S473A (96%). This data further indicates that T472 and S473 are redundant phosphorylation sites and phosphorylation at any one of these residues is a prerequisite for S450 phosphorylation. Together our data strongly establish the high specificity of ERK2 towards S450, T472 and S473 residues in NP.

### 3. ERK2 mediated NP phosphorylation is critical for virus replication

To investigate the role of ERK2 mediated NP phosphorylation in influenza virus life cycle, recombinant influenza A/WSN/1933 viruses harboring NP-S450A, NP-S473A and NP-S450A/ S473A mutations were rescued. All the mutant viruses showed lower initial titre and smaller plaque size compared to the WT virus (Fig 2A) rescued in the same experiment. The mutant viruses showed slower replication kinetics in MDCK cells (Fig 2B), with the single mutants showing almost one log attenuation compared to the WT virus by 48hpi. Viruses harboring the double mutant NP-S450A/S473A showed the most drastic two log attenuation by 48hpi. Clearly, blocking the ERK2 mediated NP phosphorylation severely restricted virus replication thereby establishing the importance of these phosphorylation events in influenza virus life cycle.

**Fig 2.**
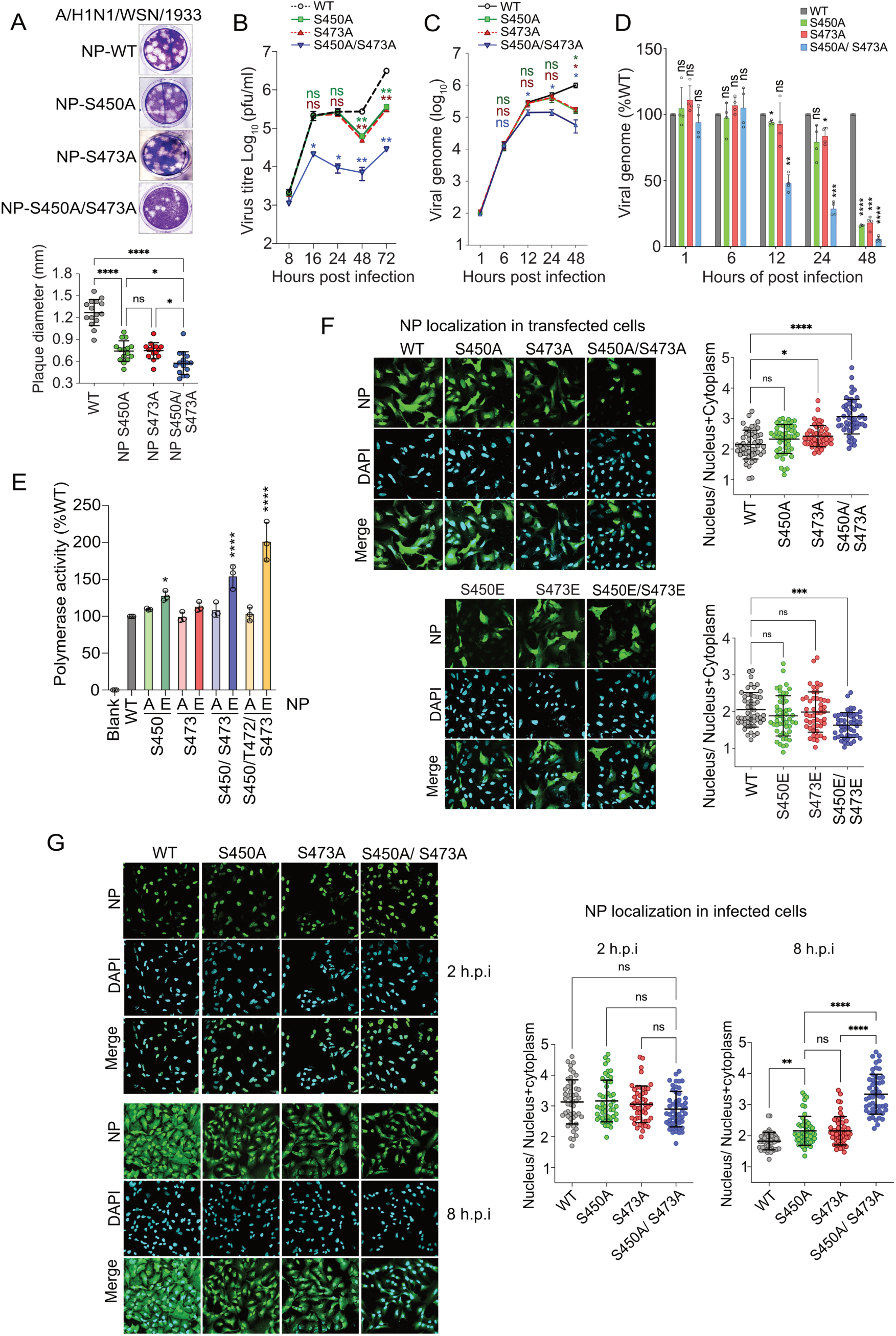
ERK2 mediated phosphorylation is critical for NP/ RNP nuclear export and virus replication. (A) Plaque morphology and average plaque diameter of WT or recombinant NP mutant viruses. (B) Multicycle replication kinetics of the mutant NP viruses. Absolute (C) and relative (D) abundance of viral genome at different times of post infection were quantified using real time qRT-PCR. (E) Polymerase activity assay using negative sense viral RNA (vRNA) template along with WT and mutant NP reconstituted in HEK293T cells. (F) WT and mutant NP proteins expressed in A549 cells were fixed,immuno-stained with anti-NP antibody at 36 h.p.t and imaged using a confocal microscope. (G) A549 cells infected with WT or mutant viruses (MOI:5). At 2 and 8 hpi cells were fixed, immuno-stained using anti-RNP antibody and imaged using a confocal microscope. 50 cells from 5 different fields were analysed using image J software to present nuclear-cytoplasmic distributions of NP.

To gain further insight regarding the role of ERK2 in modulating specific steps of influenza virus life cycle, we monitored viral RNA synthesis in cells infected either with WT or mutant viruses at different times of post infection. Quantitative RT-PCR analysis performed using viral genomic RNA (vRNA) or viral mRNA specific primers showed that wild type and mutant viruses exhibited comparable vRNA and mRNA abundance as early as 1hpi and 6hpi. However, the mutant viruses showed attenuated RNA synthesis at later time points. The double mutant virus (NP-S450A/S473A) showed 50% reduction in vRNA synthesis at 12hpi, which gradually decreased to 5% by 48 hpi (Fig 2C, D). The single mutant S450A and S473A viruses showed no or minor reduction till 24 hpi but then a drastic reduction to 16% and 18% respectively by 48 hpi (Fig 2C, D). The mRNA synthesis for WT and NP mutant viruses showed trend similar to vRNA synthesis for all the time points (Fig S6 A, B). Considering that influenza virus completes a single infection cycle by 6 hours (41), this data suggests that the mutant viruses were not defective in RNA synthesis, rather one of the post viral RNA synthesis steps was blocked. To investigate this further, reporter viral RNPs were reconstituted in HEK293T cells to monitor viral RNA synthesis independent of other steps of virus life cycle. Plasmids expressing viral reporter RNA template (either negative or positive sense), RdRp subunits and NP protein were transfected and reporter activity was monitored as a proxy of viral RNA synthesis (Fig 2E, S6C). As evident, RNPs reconstituted with phospho-null NP mutants (S450A, S473A, S450A/ S473A and S450A/ T472A/ S473A) barely showed any decrease in viral RNA synthesis, thereby ruling out the possibility of attenuation in virus replication due to defect in RNA synthesis. Interestingly, RNPs reconstituted with multiple phosphomimetic glutamic acid residue substitutions of NP (NP-S450E/ S473E and NP-S450E/ T472E/ S473E) showed higher RNP activity (Fig 2E, S6C). Both phosphonull and phosphomimetic NP mutant proteins showed expression levels comparable to the WT proteins (Fig S6D). The mutants also displayed similar trends of reporter activity for both positive and negative sense replicons (Fig 2E, S6C), thereby confirming that the NP phosphorylation is not essential for supporting viral RNA synthesis.

We have shown earlier that PKCδ mediated NP phosphorylation at its homotypic interface, regulates its oligomerization and assembly into viral RNPs (33, 36). The S450 residue is situated close to the homotypic interface of NP and participates in intermolecular interaction with the neighbouring protomer. To test whether the ERK2 mediated phosphorylation at S450, T472 and S473 also regulate NP oligomerization, we utilized recombinant, purified WT and mutant NPs and subjected them to size exclusion chromatography. As evidenced (Fig S7) WT and mutant NP proteins showed similar oligomerization propensity with minor or no changes in the relative abundance of the oligomeric and monomeric NP populations. This excludes any major role of ERK2 mediated phosphorylation in regulating NP oligomerization and RNP assembly.

### 4. ERK2 phosphorylation promotes nuclear export of NP and viral RNPs

Next, we investigated the potential role of ERK2 mediated phosphorylation in regulating NP’s nuclear-cytoplasmic distribution. Individual NP mutants harboring either phosphonull or phosphomimetic mutations at S450 and S473 residues or both were transiently overexpressed in A549 cells and their subcellular localization was monitored using immunostaining with NP specific antibody. NP showed exclusive nuclear enrichment at early times of post transfection (12-24 hours) while demonstrating a mixed nuclear-cytoplasmic redistribution at a later time point (36 hours) (Fig S8 A). NP mutants harboring phosphor-null or phosphomimetic mutants at the ERK2 phsopho-sites showed no difference in nuclear enrichment at 12 h.p.t (Fig S8B). However, NP phospho-null mutations showed low-moderate (for NP-S473A) to high (for NP-S450A/ S473A) nuclear retention compared to the WT at 36 h.p.t., hence suggesting potential role of these phosphorylation sites in regulating NP nuclear export (Fig 2F). In line with this observation, phosphomimetic substitutions both at S450 and S473 residues (NP-S450E/ S473E) renders the protein more efficient in nuclear export at 36 h.p.t thereby showing higher extent of cytoplasmic distribution with respect to the WT NP. At 24 h.p.t. both phospho-null and phosphomimetic mutants showed intermediate phenotype (Fig S8B).

In cells infected with WT A/WSN/1933 virus (MOI-5), NP showed nuclear accumulation as early as 2 hpi and mixed nuclear-cytoplasmic distribution by 8 hpi (Fig S9). In contrast, viruses harboring phospho-null NP-S450A and NP-S472A mutations showed nuclear retention of NP even at 8 hpi and this alteration in nuclear-cytoplasmic distribution was more drastic for the virus harboring the double mutant NP-S450A/ S473A (Fig 2G). These data clearly demonstrate that ERK2 mediated NP phosphorylation is critical for promoting the nuclear export of the newly assembled vRNP complexes at the late stages of infection, and this is independent of RSK1 mediated facilitation of vRNP export (22).

### 5. Active PKC-α scaffolds a stable interaction between NP, MEK1 and ERK2

To this end, we aimed to characterize the molecular interaction between ERK2 and its substrate NP, which may facilitate NP phosphorylation at specific sites mentioned above. While kinase substrate interactions could be transient, active ERK2 is known to participate in stable interaction with its substrates through recruitment of two docking domains (42). Additionally, a number of scaffold proteins mediate interactions between ERK with its substrate along with the upstream kinases like MEK and Raf (43). ERK2-NP interaction was investigated by co-expressing these two proteins followed by immunoprecipitating ERK2 and monitoring the co-immunoprecipitation of NP. No co-precipitation and hence no interaction were observed between the two proteins (Fig 3A lane 2). Considering that ERK2 activation may facilitate its interaction with NP, we included the constitutively active PKCα-CAT in this experiment, which is shown to activate ERK2 (Fig S1A) and trigger NP hyperphosphorylation (Fig 1B). Interestingly, the PKCα activated ERK2 can co-precipitate NP (Fig 3A, lane 3). Additionally, ERK2 can also co-precipitate reconstituted viral RNPs but only in the context of PKCα-CAT overexpression (Fig 3A, lane 5). Interestingly, in both cases, the hyperphosphorylated form of NP was specifically enriched through ERK2 co-precipitation, hence suggesting that ERK2-NP interaction facilitates NP/ RNP phosphorylation. Apart from WSN NP, the A/H1N1/PR8 and B/Brisbane NP, which were shown to get phosphorylated by ERK2 (Fig1D), also co-precipitated with ERK2 in presence of PKCα-CAT (Fig 3B & 3C), thereby suggesting conserved nature of this kinase-substrate interaction in different influenza viruses. To test if MEK, the upstream kinase of ERK2, can also interact with NP, co-immunoprecipitation assay was performed with MEK1 or MEK2 co-expressed with NP either in presence or absence of the PKCα-CAT. Similar to ERK2, MEK1 but not MEK2 interacted with NP in a PKCα dependent manner (Fig 3D). Conversely, NP can also co-precipitate ERK2 and MEK1 in the reverse-coimmunoprecipitation assay, but only in the presence of PKCα-CAT and the extent of co-precipitation of MEK1 is much higher than that of ERK2 (Fig 3E).

**Fig 3.**
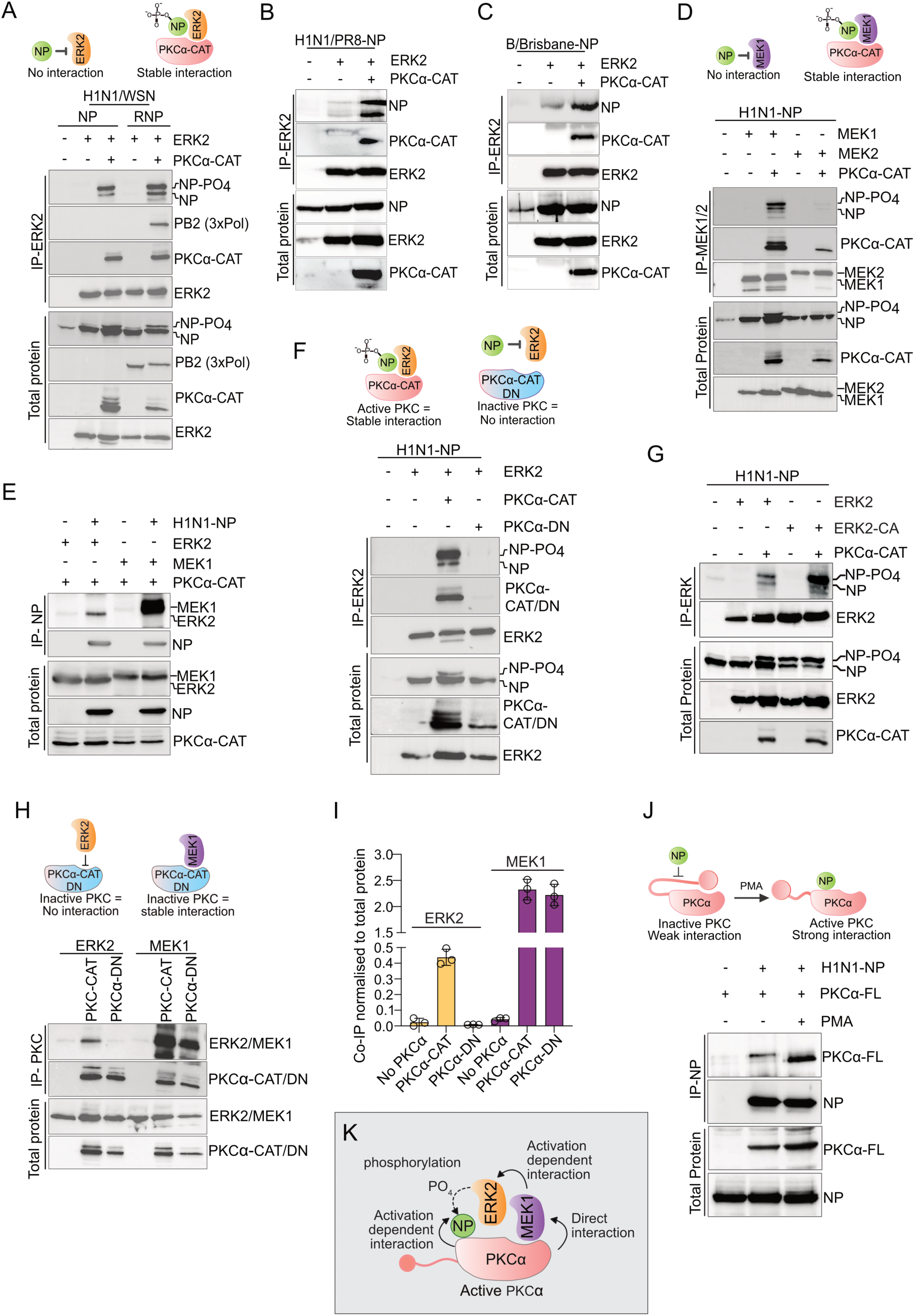
PKC-α scaffolds a stable interaction between viral NP/ RNP with MEK1 and ERK2. (A) Co-immunoprecipitation of H1N1/WSN NP/ RNP with ERK2. Cells were transiently transfected to overexpress NP-V5 or to reconstitute complete RNP complexes either in the context of ERK2-FLAG or PKCα-CAT-HA overexpression and immunoprecipitated using ERK2 specific antibody.(B) Co-immunoprecipitation of A/H1N1/PR8 NP and (C) B/Brisbane NP with ERK2. (D) Co-IP of NP with MEK1/2-FLAG in cells overexpressing NP either with MEK1/2 or MEK1/2 and PKCα-CAT. (E) Reverse Co-IP showing co-precipitation of ERK2-FLAG and MEK1-FLAG with immunoprecipitated NP. (F) Co-IP of NP-V5 with ERK2-FLAG either in the context of PKCα-CAT-HA or PKC-α-DN-HA overexpression. (G) Co-IP of NP with ERK2-FLAG or constitutively active ERK2-FLAG (ERK2-CA) in presence or absence of PKC-α-CAT-HA overexpression. (H) Co-IP of ERK2-FLAG and MEK1-FLAG with PKCα-CAT-HA and PKCα-DN-HA. (I) Densitometric analysis of Fig H. (J) Co-IP of full length PKCα (PKCα-FL) with NP overexpressed in cells stimulated with PMA or with vehicle control. (K) Model describing the scaffolding role of active PKCα. All experiments are performed with A/H1N1/WSN/NP except otherwise stated.

PKCδ is known to interact with NP/ RNPs through the intermediacy of PB2 subunit of the viral RdRp (33). Considering its close association with viral RNPs, we asked if PKCδ can also mediate NP’s interaction with ERK2. ERK2-NP co-immunoprecipitation was performed in presence or absence of constitutively active catalytic domains of this PKC isoforms. We observed that PKCα-CAT alone and not PKCδ-CAT facilitated coprecipitation of NP with ERK2 (Fig S10 A), thereby establishing the exclusivity of PKCα in mediating this interaction. Interestingly, a catalytically inactive version of PKCα-CAT (designated here as the dominant negative: PKCα-DN; reason described later) failed to facilitate NP coprecipitation with ERK2 (Fig3F lane 4), hence suggesting that active PKCα is essential for supporting ERK2-NP/ RNP complex formation (Fig 3F).

There could be two different models supporting PKCα mediated NP’s interaction with MEK1 and ERK2. First, PKCα can activate the MEK1/ ERK2 cascade and hence facilitate ERK2 mediated NP phosphorylation. Second, PKCα may directly interact with MEK1, ERK2 and NP to act as a scaffold protein, thereby leading to the robust activation and high specificity of the MAPK cascade towards its substrate, NP. In order to test these hypotheses, we generated a constitutively active ERK2 (ERK2-CA) harboring L75P/S153D/D321N mutation (44) and tested its ability to interact with NP in presence and absence of PKCα (Fig 3G). Although ERK2-CA triggered robust NP hyperphosphorylation, but failed to directly interact with NP in absence of the PKCα. However, in presence of PKCα-CAT, ERK2-CA showed higher extent of NP co-precipitation compared to the WT ERK2. This data suggested that PKCα-CAT not only activates the MEK1-ERK2 cascade but also physically bridges the interaction of these kinases with NP. To validate this model further, interaction ability of PKCα-CAT along with MEK1, ERK2 and NP was investigated using co-immunoprecipitation analysis. Surprisingly, PKCα-CAT co-precipitated both ERK2 and MEK1 (Fig 3H & I), and the extent of co-precipitation was 5 fold higher for the latter one. Thus, active PKCα showed stronger affinity for MEK1 than ERK2. This explains why MEK1 showed higher co-precipitation with NP than ERK2 in presence of PKCα-CAT (Fig 3E). Interestingly, the catalytically inactive PKCα (PKCα-DN) failed to co-precipitate ERK2 but retained its ability to interact with MEK1 (Fig 3H), thereby explaining why active PKCα is indispensable for the mediating ERK2-NP interaction (Fig 3A). PKCα-CAT also interacted with NP when co-expressed and unlike PKC-δ, this interaction is independent of viral PB2 protein (Fig S10;(33).

Finally, we checked if the full length PKCα (PKCα-FL) could interact with influenza virus NP protein. As shown (Fig 3J) PKCα-FL, which usually remain in the inactive conformation (45), was able to co-precipitate low levels of NP but the co-precipitation was significantly increased upon PMA stimulation. Together, our data show that PKCα, in its active form, physically interacts with NP, MEK1 and ERK2 to form a multi-kinase complex (Fig 3K) which is responsible for ERK2 mediated direct phosphorylation of NP. This also established PKCα as a scaffold protein for the MEK/ERK pathway.

### 6. The NP associated multi-kinase complex shows a dynamic nuclear-cytoplasmic accumulation at different stages of infection

To this end we aimed to gather evidence in support of the formation of the stable complex between the host kinases and NP during the course of infection. In this regard, A549 cells were infected either with A/H1N1/WSN and B/ Brisbane/60/2008 (MOI-10) viruses, respectively. NP/RNP complexes were immunoprecipitated at 6 hpi and co-precipitation of endogenous PKCα and ERK2 was assessed using specific antibodies (Fig 4A, B). Interestingly, both PKCα and ERK2 were co-precipitated with NP/RNPs for both types of influenza viruses. Interestingly, the extent of co-precipitation was higher for PKCα than that of ERK2, thereby supporting the key role of PKCα in bridging the NP-ERK2 interaction. This data along with the results presented in figure 3, confirms the existence of the multi-kinase complex, constituted of PKCα, MEK1 and ERK2, which stably associates with different influenza virus RNPs during infection.

**FIG 4.**
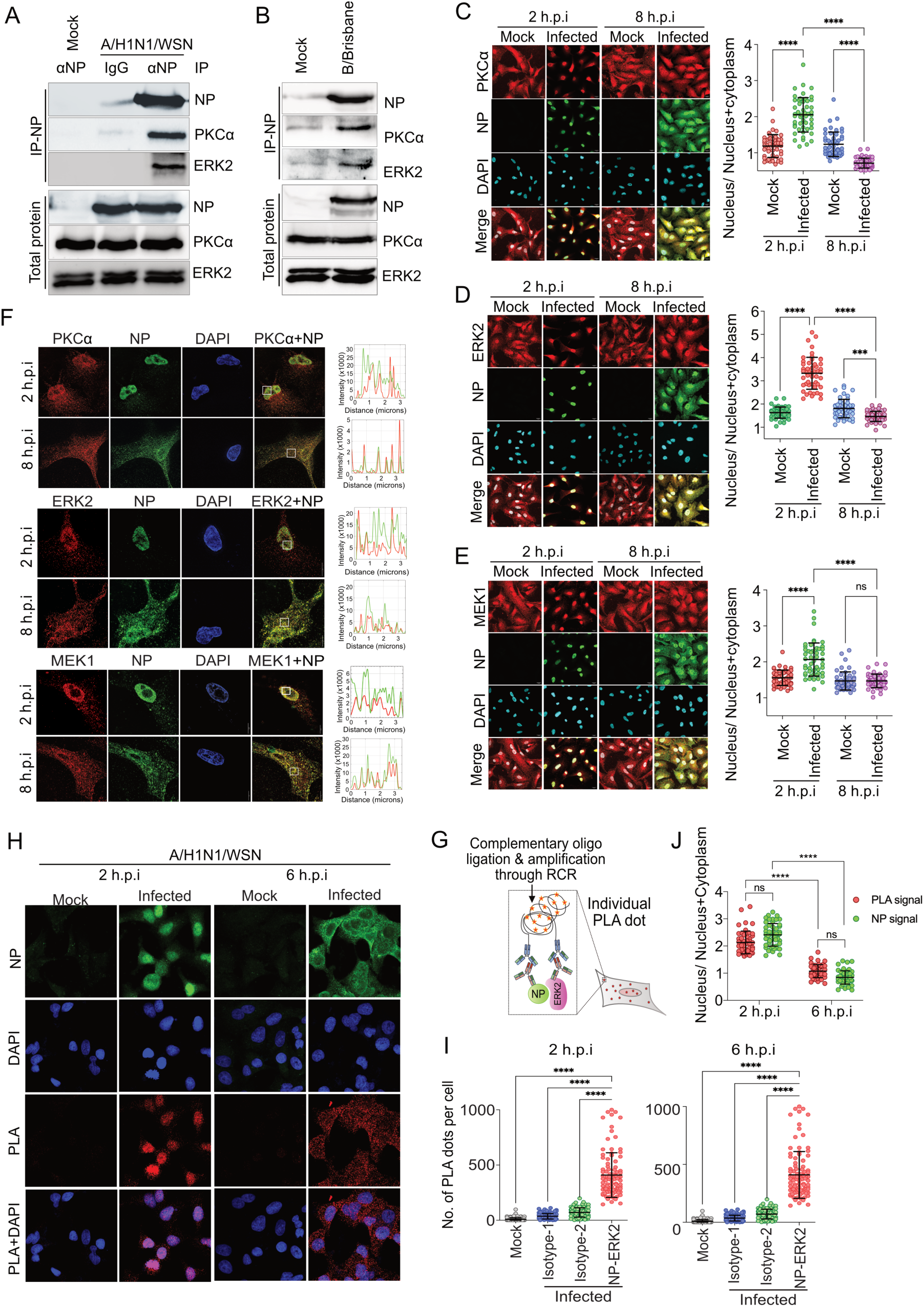
The NP associated multi kinase complex formation and its dynamic localization in virus infected cells. (A) Immunoprecipitation of NP from A/H1N1/WSN virus infected A549 cells (MOI-10) at 6 hpi using ANP antibody. (B) B/ Brisbane/ 60/ 2008 virus infected (MOI-10, 6 hpi) A549 cells subjected to immunoprecipitation using antiBNP antibody. A549 cells infected with influenza A/ H1N1 virus (MOI-5) were fixed at 2 and 8 hours of post infection and immunostained for NP and PKCα (C) or ERK (D) or MEK (E). Images were analysed with image J software to present subcellular distribution of the proteins (lower panel). (F) Super-resolution micrographs of the infected A549 cells showing precise co-localization of viral NP with PKCα, ERK and MEK at 2 and 8 hpi. Co-localization of PKC, ERK and MEK (in red line) with NP (in green line) was plotted through Image J software. (G) Schematic representation of proximity ligation assay. (H) Cells infected with influenza A/H1N1 virus or mock infected were subjected to PLA using anti-RNP (goat) and anti-ERK2 (rabbit) primary antibodies or with one of the primary antibody along with the IgG isotype control for the other (full panel shown in Figure S11). Post PLA, cells are stained with anti-NP antibody (mouse) and imaged. (I) Images were analysed for the PLA dots per cell using Image J software and plotted for quantitative representation. (J) Subcellular distribution of PLA dots and NP were analysed using image J software.

Components of the MEK-ERK cascade localizes within the cytoplasm in the resting cells but can be imported into the nucleus upon extracellular stimulations (22, 28, 35). We checked if the subcellular distribution of endogenous ERK2, MEK1 and PKCα coincide with the dynamic nuclear-cytoplasmic localization of NP during the course of infection (Fig S9). A/WSN/1933 infected A549 cells showed significant nuclear enrichment of PKCα, ERK2 and MEK1 compared to the mock infected cells at 2 hpi but not at 8 hpi (FIG 4 C-E). Clearly, PKCα along with MEK and ERK colocalize with NP within the nucleus early during infection. To further investigate this, we performed super-resolution microscopy of the infected A549 cells, and observed punctate structures of NP/ RNP complexes in the nucleus at 2 hpi and both in nucleus and cytoplasm at 8 hpi. At both the time points, ERK, MEK and PKCα proteins individually showed high co-efficient of colocalization with the punctuated structures of influenza NP/ RNP complexes at a close proximity of 50-60 nM approximately. Additionally, majority of the co-localization was observed within the nucleus early during infection and in the cytoplasm during the late stage (Fig 4 F, Table S8). These data suggest that the localization of ERK, MEK and PKCα dynamically changes with NP during the course of infection resulting in close association between these host proteins with viral RNPs, thereby facilitating interaction between them.

We probed the NP-ERK2 interaction to analyse the spatiotemporal dynamics of the RNP associated multi-kinase complex formation during the course of influenza virus infection. We performed proximity ligation assay (PLA), wherein close association (within 40nm) of the partner protein molecules appears as a fluorescent foci within the cell and the number of the foci in each cell could serve as a quantitative estimate of the corresponding protein-protein interaction (Fig 4G). A549 cells were either mock infected or infected with A/WSN/1933 virus, fixed at 2 hpi and 6 hpi and were subsequently subjected to PLA followed by detection of the fluorescent signal. Localization of viral NP was visualised using immunostaining post PLA treatment. Highly abundant fluorescent PLA dots were detected in the infected cells incubated with both anti-ERK2 rabbit and anti-RNP goat (majorly detects NP) antibodies but no signal was obtained for the mock infected cells or the infected IgG isotype treated control cells (Fig 4H, I, S11). This further confirmed a stable association between ERK2 and NP/RNP during the course of infection. Interestingly, the PLA dots were majorly located within the nucleus at 2 hpi but were distributed throughout the cytoplasm at 6 hpi, which perfectly coincides with the localization of NP within these virus infected cells (Fig 4J). Thus, the PLA result along with data presented so far collectively suggested that the ERK2-NP complex formation happened within the nucleus, through the intermediacy of PKCα and MEK1, at early times of post infection. The same complex however, might get exported out to the cytoplasm at late hours (6-8 hpi), possibly mediating the export of the newly assembled vRNP complexes.

### 7. PKCα serves as the key regulator of vRNP export and influenza virus replication

To this end, we aimed to establish that PKCα mediated ERK2-NP complex formation is critical for vRNP export and hence production of infectious virus particles. We used the catalytically inactive PKCα-DN, which can interact with MEK1 but failed to interact with ERK2 (Fig 3H) and hence unable to bridge the ERK2-NP interaction (Fig 3F). We postulated that PKCα-DN, upon overexpression in cells, should sequester out endogenous MEK1 and interfere with the PKCα-MEK1-ERK2-NP complex formation. Thus, PKCα-DN should act as dominant negative over the endogenous PKCα in terms of promoting ERK2 mediated NP phosphorylation and supporting the vRNP export at later times of infection. To validate this hypothesis, A549 cells overexpressing PKCα-DN were infected with influenza A/WSN/1933 and the NP/ RNP localization was monitored at 2 and 8 hpi (Fig 5A). As postulated, over expression of PKCα-DN severely restricted NP within the nucleus at 8hpi. Interestingly, the number of NP positive cells was also reduced more than 50% with respect to the control set (Fig 5B), which indicated significant retardation of virus replication as early as 8hpi. This reduction became even more prominent during multicycle replication where PKCα-DN overexpression resulted in two log reduction in virus titre at 12-36 hpi (Fig 5C). This established the critical role of PKCα in influenza A virus RNP export and virus replication, through mediating a stable ERK2-NP interaction and thereby promoting ERK2 mediated NP phosphorylation.

**Fig 5.**
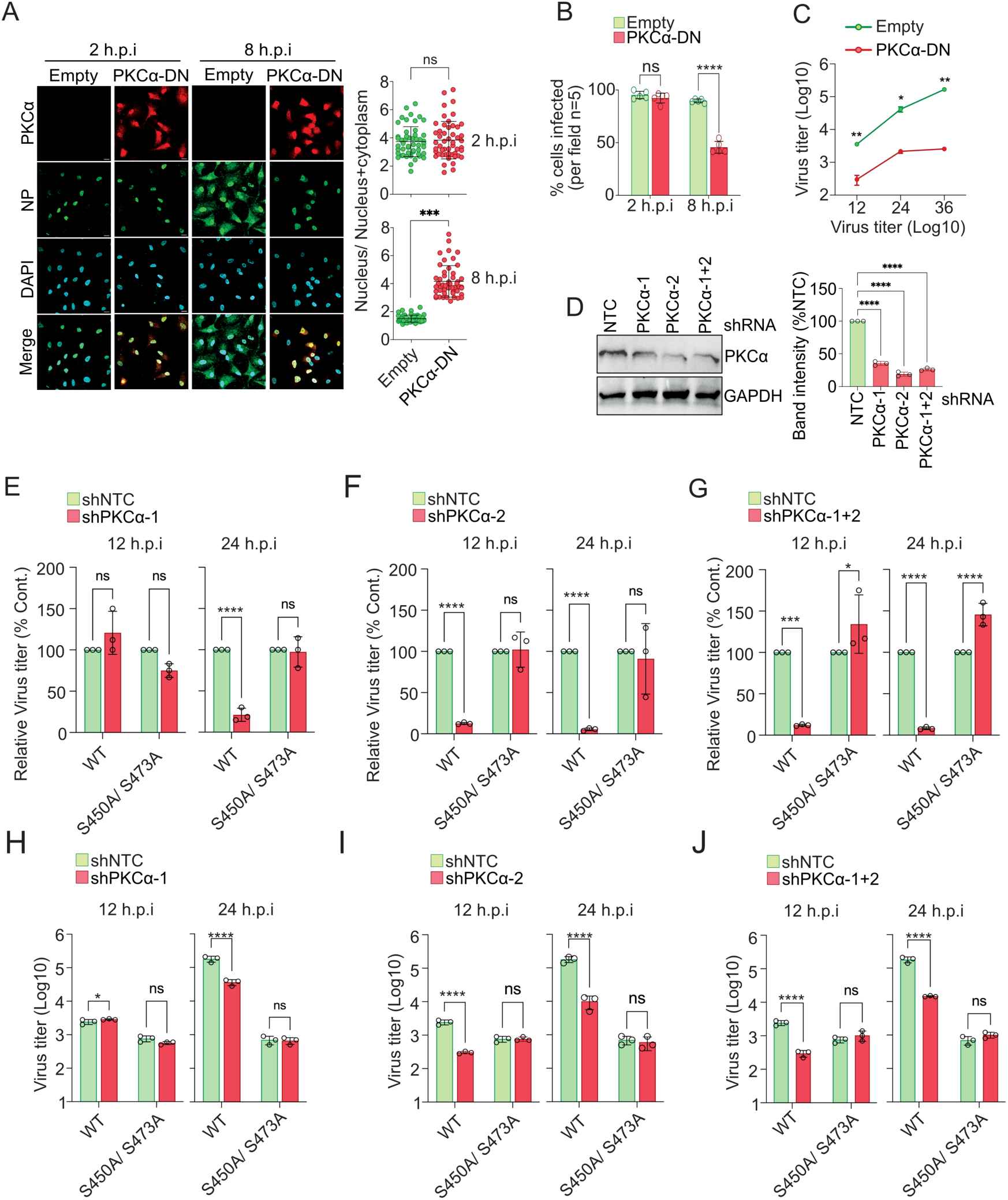
PKCα as the key modulator of influenza virus replication. (A) Cells overexpressing PKCα-CAT-DN or control cells were infected with A/H1N1/WSN/1933 virus followed by fixing and immunostaining at indicated times of post infection. (B) Images from five different fields were analyzed to measure the extent of infection using image J software. (C) Virus titers from PKCα-CAT-DN overexpressing or control cells were measured using plaque assay at different times of post infections as indicated. (D) A549 cells were transduced with lentiviruses expressing PKCα specific or non-target (NTC) shRNAs. Knockdown efficiency was tested by western blot analysis and PKCα band intensity from three different experiments are plotted using image J software. PKCα deficient cells (PKCα-KD1, PKCα-KD2, PKCα-KD1+2) were infected with WT or NP-S450A/ S473A mutant viruses and titer was measured at 12 and 24 hpi. Relative (D-F) and absolute (G-I) virus titers for both the viruses at two different time points are plotted.

To draw a more direct correlation between PKCα’s role in promoting virus replication and ERK2 mediated NP phosphorylation, we generated PKCα deficient stable cell lines using shRNA mediated knockdown. Three different cell lines expressing shPKCα-1, shPKCα-2 and shPKCα-1+2 showed varied degrees of knockdown with the latter two achieving around 80% reduction in the PKCα level compared to the control cells (shNTC) (Fig 5D). A remarkable reduction in viral titer was observed in shPKCα-1 cells at 24 hpi and also in shPKCα-2 and shPKCα-1+2 cells at both 12 and 24 hpi, compared to target shNTC cell line (Fig 5, E-J). Interestingly, the double mutant NP-S450A/ S473A virus showed similar extent of replication in both the control and knockdown cells at both the times of post infection (Fig H-J). Clearly, disruption of the ERK2 phosphorylation sites made the NP (S450A/S473A) mutant virus insensitive towards the depletion in active PKCα abundance in cell. This data unambiguously established the critical role of PKCα in regulating ERK2 mediated NP phosphorylation, thereby promoting vRNP export and ultimately facilitating the production of infectious virus particles.

## Discussion

Influenza virus RNP’s optimal life span within the host cell nucleus and their timely export to the cytoplasm is essential for viral gene expression and genome replication and production of infecctious virus particles. This timely coordinated nuclear-cytoplasmic shuttling of influenza virus RNPs is tightly regulated through host kinase mediated phosphorylation of NP at multiple serine, threonine and tyrosine residues, and ERK activated Rsk1 has been identified as one such kinase to be involved in vRNPs nuclear export (18–22). In this study, we show for the first time that ERK2 can directly phosphorylate NP at distinct serine and threonine residues which promote vRNP’s flight from nucleus to cytoplasm. In addition, our study unravels a unique mechanism of PKCα mediated activation of the MEK-ERK pathway. We show that active PKCα scaffolds a stable tripartite interaction between MEK1, ERK2 and viral NP/ RNPs. This RNP associated multi-kinase complex is formed within the nucleus during early phase to promote NP phosphorylation and dynamically translocate to cytoplasm at later phase of infection.

Using Analogue sensitive kinase, we established a direct kinase-substrate relationship between host ERK2 and influenza virus NP. ERK2 phosphorylates A/H1N1/WSN NP at Ser450, Thr472 and Ser473 residues wherein the later two residues are situated within a consensus “Thr472-Ser473-Pro474” motif (46), a putative ERK2 phosphorylation site. Our data showed that phosphorylation at one of the consensus Ser/Thr residues is pre-requisite for the phosphorylation at the non-consensus Ser450 residue. Interestingly, S450 and T472 phosphosites remain conserved across different influenza viruses suggestive of their essential role in supporting virus replication cycle. In line with this observation, mutation of two out of the three phospho-acceptor sites (S450A/ S473A) makes the virus severely attenuated compared to the WT virus.

The MAPK, MEK-ERK pathway has long been implicated in RNP nuclear export (25, 28). Corroborating with this notion, phospho-null or phosphomimetic substitutions at the ERK2 phosphorylation sites imparted a negative or positive influence upon the RNP nuclear export. Additionally, phospho-mimetic substitutions at the phosphorylation sites boosted viral RNA synthesis, while the phosphonull substitutions failed to show any significant effect. The mutant viruses harboring phospho-null substitutions showed no defect in RNA synthesis during the early cycles of replication, rather demonstrated limited vRNP export and reduced virus titre, which indirectly impacted viral RNA synthesis at later infection cycles. These data suggest that ERK2 mediated NP phosphorylation is essential for vRNP’s transport from nucleus to the cytoplasm. It is noteworthy that, both ERK2 and its downstream kinase RSK1 phosphorylates distinct residues in NP to promote vRNP nuclear export (22), hence play functionally redundant roles in virus life cycle. However, it is not surprising that influenza viruses recruit multiple kinases within the MAPK cascade to tightly regulate vRNP nuclear export thus ensuring optimal virus growth.

ERK2’s specificity towards its substrate can be determined through several factors including (i) cross-talk with other signalling cascades (47), (ii) dynamic alteration in subcellular localization (48) and (iii) interaction with scaffold proteins (49). Our data suggests that influenza virus employs all of these strategies for ensuring ERK2 mediated specific phosphorylation of NP/ RNP complexes. First, we showed that PKCα, which is not an integral part of the MAPK cascade, activates ERK2 to phosphorylate NP (Fig 1). Second, influenza virus infection triggers PKCα, MEK1 and ERK2 translocation to the nucleus as early as 2hpi, coordinating with the localization of the newly synthesized NP proteins to the nucleus (Fig 4). Third, we present multiple evidences highlighting that PKCα acts as a scaffold to form a stable NP-PKC-MEK1-ERK2 complex (Fig 3, 5). Interestingly, PKCα shows higher affinity towards MEK1 than ERK2 and the interaction with MEK1 is independent of its activation status. This is suggestive of a model where active PKCα interact with NP and MEK1 directly, leading to MEK phosphorylation and subsequent activation. The PKC bound active MEK1 then interacts with ERK2 leading to its activation and also bringing it to the close proximity to its substrate, NP. In this regard it should be noted that the MEK1 is known to interact with ERK2, which helps ERK2 to transition to a conformation that is ideal for interaction with its substrate (50). Thus, the scaffolding activity of PKCα creates a platform where the effector kinase ERK2 remains in close association with its activator kinase MEK1 and its substrate NP, thereby ensuring selective activation and high signaling specificity of the MAPK cascade. This model is indirectly supported by the fact that the catalytically inactive PKCα (PKCα-DN), which can co-precipitate MEK1 but not ERK2, failed to bridge NP-ERK2 interaction. In fact, the PKCα-DN, when over-expressed during the course of infection, imparts a dominant negative phenotype, blocking the vRNP export and perturbing virus replication. This is further substantiated by the fact that PKCα knockdown significantly impacted replication of WT influenza virus but did not alter the replication of the mutant virus lacking the ERK2 phosphorylation sites in NP. Clearly, PKCα serves as a key regulator of influenza virus life cycle (i) by activating the MAPK pathway, (ii) mediating a stable interaction between different members of the MAPK cascade and influenza virus NP/ RNPs and (iii) triggering ERK2 mediated NP phosphorylation which ultimately promote RNP nuclear export and progeny virus production.

MAPK mediated NP phosphorylation is known to regulate the vRNPs association with M1 and indirectly with CRM1, thereby promote the assembly of vRNP-nuclear export complex (18–22). However, whether these phosphorylation occur prior to, during, or post RNP synthesis, is not clear yet. Our immunofluorescence data suggests that activated PKCα, MEK1 and ERK2 translocate to the nucleus and co-localizes with NP as early as 2hpi. PLA data also showed that ERK2 stably associate with NP during the same time within the nucleus and together gets exported out to the cytoplasm at later time points (8hpi). Possibly, the newly synthesized NP molecules are quickly phosphorylated by ERK2 post nuclear translocation before participating in the RNP assembly process. The ERK2 phosphorylated NPs may serve as superior substrates for RNP assembly, as indicated by our RNP activity assay. Once assembled, these phosphorylated RNPs gets exported out to the cytoplasm through CRM1 dependent or independent pathways. This might be reason why the NP mutant viruses lacking the Rsk1 phosphorylation sites replicated to a significant extent (22), facilitated by ERK2 mediated phosphorylation. The existence of ERK2-NP/ RNP complexes at the cytoplasm at the later stages of infection suggests that this RNP associated host kinase complex may also regulate post export trafficking and virion assembly process which warrants further investigation.

In summary, we elucidate a novel mechanism highlighting the role of PKCα mediated activation of MEK-ERK pathway which involve formation of a multi-kinase complex that associate with influenza virus NP protein and phosphorylates it to promote vRNP nuclear export and production of infectious virion particles. The scaffolding activity of PKCα in mediating a MAPK-substrate interaction has never been reported for any other physiological processes, which suggests that influenza viruses utilize novel pathways to exploit multiple host kinases to facilitate its life cycle. Characterizing these interactions and understanding their spatiotemporal dynamics is critical for the detail understanding of the virus-host kinase relationship. This will open up new scopes to target such pivotal virus-host interactions for developing new antiviral strategies.

## Materials and Methods

A549 cells were infected with either A/WSN/1933 (H1N1) or influenza B (B/ Brisbane/ 60/ 2008) viruses at respective MOI for 1h at 37°C for indicated time points followed by immunoprecipitation using NP specific antibodies to isolate the multi-kinase complexes. For multicycle kinetics assay, A549 cells infected with either A/WSN/1933 (H1N1) WT or recombinant phosphonull mutant NP viruses. Alternatively, A549 cells stably expressing either shPKCα or shPKCαKD(1+2) or transiently over expressing pcDNA3 or PKCα-DN were infected with either A/WSN/1933 (H1N1) WT or mutant NP recombinant viruses for indicated time points. Supernatant were collected and progeny viral titer was determined by plaque assay.

Influenza virus or host cellular proteins were transiently expressed in HEK293T cells using Lipofectamine 3000 (Invitrogen). Cells were lysed in radio-immunoprecipitation assay (RIPA) buffer, immunocaptured using respective antibodies on specific beads as mentioned. Immunoprecipitated proteins were analyzed by western blotting.

Invitro kinase assay was performed by purifying active FLAG tagged ERK2, MEK1, MEK2 proteins overexpressed in HEK 293T cells and stimulated with PMA. Protein-A magnetic sure beads (Biorad) containing immuno-captured kinases were incubated with WT or mutant recombinant viral NP proteins (pretreated with RNaseA for 1 h) in the presence of γ-32P ATP at 30°C for 45min followed by autoradiography. For identification of the phospho-sites in vitro kinasing product was subjected to or LC-MS/MS analysis.

Analogue sensitive ERK2 (Q105G) immuno-purified from HEK293T cells was subjected to in vitro kinase assay with recombinant NP substrate using thio-ATP-analogue (N6-Phenylethyl γ-thio-ATP, Biolog Life Science Institute) or thio-ATP followed by alkylation using 2.5 mM of p-nitro benzyl mesylate (PNBM) at 22^0^C for 2h. Thio-phosphorylated substrates were detected by western blot using anti-thiophosphate ester antibody [51-8] (abcam).

In cell kinasing assay was performed by infecting HEK 293T cells overexpressing ERK2-AS or empty vector with Flu A/WSN virus. Cells were stimulated with PMA, or left untreated, permeabilized with 50 ug/ml digitonin (Sigma) followed by in cell kinasing with 100uM N6-phenethyl ATP-γ-S. Cells were lysed to release incell kinased NP substrate, followed by immuno capture of NP, alkylation and analysis by immunoblotting.

A549 cells grown on eight well chamber glass slides were infected with influenza A virus as mentioned. PLA was performed according to the manufacturer’s instructions. The cells were probed with primary ERK2 antibody raised in rabbit (Cell signaling) and RNP antibody raised in goat (BEI resources) at 37^0^C for 1h. PLA signals were detected at 640nm wavelength as distinct fluorescent dots or puncta using Olympus FV3000 confocal microscope. Detailed material and methods are described in SI Appendix, Material and Method.

## Acknowledgement

A.M. acknowledges the Core Research Grant (CRG) from Anusandhan National Research Foundation (ANRF)/ Science and Engineering Research Board (SERB-DST (CRG/2022/003628) for providing financial support. AR acknowledges support from Institutional Research Grant IRG-21-145-25 from the American Cancer Society (ACS). I.D.J. acknowledges ANRF/ SERB for the National PostDoctoral Fellowship (PDF/2023/000284). S.D. acknowledges the support of an MHRD for PMRF doctoral fellowship. M.S. acknowledges U.G.C for his fellowship. We acknowledge the Confocal Microscope facility supported by DST-FIST grant from Department of Bioscience and Biotechnology, Govt. of India; File no. SR/FST/LS-I/2019/595(C). We also thank Ms. Elizabeth Remily-Wood, Director of the USF MCOM Proteomics facility. The proteomics work has been supported, in part, by the Morsani College of Medicine at the University of South Florida.

## Supplementary Materials and Methods

### Cells, viruses, drugs and antibodies

Human lung epithelial cells (A549) (CCL-185), Human embryonic kidney (HEK) 293T cells (CRL-3216) and Madin Darby Canine Kidney (MDCK) (CCL-34) cells were maintained in Dulbecco’s modified Eagle’s medium (DMEM) supplemented with 10% FBS at 37°C and 5% CO_2_ along with antimycotic, penicillin and streptomycin antibiotics (Gibco).

Influenza A virus strains, A/WSN/1933 (H1N1) encoding PB2 with a C-terminal Flag (WSN-PB2-FLAG) (1) and influenza B (B/Brisbane/60/2008) viruses were used for infection in different cell lines respectively.

Antibodies used include anti-thiophosphate ester antibody [51-8] (Abcam), anti HA(C29F4,CST), anti-V5 (D3H8Q, CST), anti-NP (H16-L10-4R5) (BEI Resources), anti-FLAG M2 (Sigma), anti-influenza virus RNP (BEI Resources NR-3133), MEK1/2 (L38C12, CST) and p44/42 MAPK (ERK1/2) (137F5, CST), PKCα (#2056, CST), phospho ERK2 (CST) and, GAPDH (Sigma).

Phorbol 12-myristate 13-acetate (PMA) (Sigma) was reconstituted in DMSO at a stock concentration of 100ug/ml. As indicated in the experiments, cells were stimulated with 1.5ug/ml of PMA for 1 h or mentioned otherwise.

### Plasmids

All viruses related genes were derived from either influenza A (A/WSN/33) or influenza B (B/Brisbane/60/2008) viruses. Plasmids encoding WSN viral NP, pCDNA6.2-NP-V5 and polymerase proteins-pCDNA3-PB2-3xFLAG (encoding a C-terminal FLAG tag), pCDNA3-PA and pCDNA3-PB1 were expressed in HEK 293T cells. vNA-luc reporter plasmids encoding firefly luciferase in the negative sense flanked by UTRs from the NA gene was utilized (Regan et al., 2006). Plasmids expressing full-length isoform of PKCα (PKCα-FL, Addgene plasmids #21232), the catalytic domain (PKCα-CAT) (Addgene plasmids #21234) (2) and PKCα-CAT-DN (with K368R mutation in the PKCα-CAT construct was used to generate catalytically inactive form (3). PKCδ-CAT was obtained from (Addgene plasmids #16388). pET28a-NΔ7NP was used for bacterial expression of WT ANP with a C-terminal His tag and seven amino acid deletion on the N-terminus, as described previously (4). pET28aBNP was used for expressing BNP in bacteria. The pcDNA-ERK2-3xFLAG, the pcDNA-MEK1-3xFLAG and the pcDNA-3xFLAG constructs were generated in the lab by isolating total RNA from A549 cells, followed by RT-PCR and further amplification of specific gene of interest using gene specific primers. Constitutively active form of ERK2 was generated by introducing L75P/S153D/D321N mutation in pcDNA3-ERK2-3XFLAG plasmid (5). All mutations thus generated were confirmed by Sangers sequencing.

### Generation of influenza A virus mutant NP plasmid constructs by Site Directed Mutagenesis (SDM)

Mammalian expression vector of NP protein (pCDNA6.2-NP-V5) and bacterial expression vector-pET28a-NΔ7NP-E339A expressing monomeric NP protein, from influenza A virus was subjected to site-directed mutagenesis to generate S450A, T472A, S473A, S450A/ S473A, T472A/S473A and S450A/T472A/S473A mutations, using the QuikChange II Site-directed mutagenesis kit (Agilent Technologies) according to the manufacturer’s instructions, further verified using Sangers DNA sequencing.

### PKCα knock down cell line generation

PKCα knock down cell line were generated by using mission shRNA (Sigma) specifically targetting *PRKCA* gene in A549 cells. Inorder to generate *PRKCA* gene knockdown lentivirus, we have transfected HEK293T cells with plasmid containing either (a) pLKO.1-puro shRNAs (TRCN0000001690) targeting *PRKCA* (CTTTGGAGTTTCGGAGCTGAT) (shPKCα-1) or (b) pLKO.1-puro shRNAs (TRCN0000001691) targeting *PRKCA* (CGAGCTATTTCAGTCTATCAT) (shPKCα-2) or (c) (shPKCα-1+2) together or (d) a non-targeting control pLKO.1-puro non-mammalian shRNA control plasmid DNA (NTC), SHC002 plasmids along with psPAX2 (Addgene #12260) and pVSVG (Addgene#138479) till 48hrs. Lentiviral particles thus produced were either used alone or in combination to transduce A549 cells. Transduced cells shPKCα-1, shPKCα-2 and shPKCα-1+2 thus produced were subjected to puromycin (5ug/ml) selection. *PRKCA* gene knockdown was confirmed by western blot analysis of the cell lysate. All experiments using the knockdown cell lines were performed with early passages of knockdown cell lines.

### Generation of recombinant NP mutant viruses

The mutations S450A, S473A and S450A/S473A were introduced in to the bidirectional pBD-NP plasmid DNA by SDM. Rescue of recombinant viruses was performed as described previously by Hoffmann et al., 2000 (6). Briefly, 1.5ug pTMΔRNP plasmid (kind gift from Andrew Mehle) along with 0.25ug of pBD-PB2, pBD*-PB1, pBD-PA and pBD-NP (WT or having desired mutations) plasmids were transfected in 1.5× 10^6^ HEK293T and 0.5× 10^6^ MDCK cells using Lipofectamine 3000 according to the manufactureŕs protocol. 24 h post transfection the initial medium was changed to virus growth medium (VGM)(3) supplemented with 0.5 µg/ml TPCK-treated trypsin and the supernatant (P0) was harvested at 48 h.p.t. Viral titers were determined by standard plaque assay. Subsequently amplified through two consecutive passaging and the cDNA of the P2 virus was subjected to Sangers sequencing. The P2 virus was utilized for subsequent characterization experiments.

### Multicycle replication assay

Influenza A virus multicycle replication kinetics assays were performed by following the protocol used previously by Mondal et al 2017 (3). A549 cells stably expressing either shPKCα or PKCα-Knock down stable cell were infected with either A/WSN/1933 (H1N1) WT or mutant NP viruses (S450A, S473A, S450A/S473A respectively) at a MOI of 0.01.

A549 cells transiently expressing pcDNA3 or PKCα-DN were subsequently infected with A/WSN/1933 (H1N1) viruses at an MOI of 0.01. Supernatant containing virus particles were collected at different time points and subsequently viral titers were determined by performing plaque assay on MDCK cells (3).

### Invitro kinase assay

The Invitro kinase assay was performed according to the protocol described by Ludwig et al 1996 (7). Briefly, HEK 293T cells were transfected with FLAG tagged ERK2, MEK1, MEK2 plasmids for 23hrs. The cells were left unstimulated or stimulated with PMA for 1hr followed by lysis in modified RIPA buffer (25 mM Tris [pH 7.5], 137 mM NaCl, 0.1% SDS, 0.5% Deoxycholate, 1% NP 40, 2mM EDTA, 10% glycerol, protease and phosphatase inhibitors) and immunoprecipitation using FLAG antibody. Immune complexes were captured with Protein A magnetic beads, which were then washed once in Triton lysis buffer (TLB; 20 mM Tris–HCl pH 7.4, 500 mM NaCl, 10% glycerol, 1% Triton X-100, 2 mM EDTA, protease and phosphatase inhibitor) and then twice in ERK2 kinase buffer (MgCl_2_ 10mM, 25mM HEPES pH 7.5, 0.5mM DTT, protease and phosphatase inhibitors) and finally resuspended in the same buffer. Wild type (Wt) and mutant recombinant viral NP proteins were purified from bacteria following the existing protocol and treated with RNaseA for 1 hr prior to be use as substrate (4). 4ug of the NP protein was incubated with protein-A magnetic sure beads (Biorad) containing immuno-captured kinases in the presence of 10 mCi of γ-32P ATP and 5mM of cold ATP at 30°C for 45min with intermittent shaking every 10 minutes. Reactions were terminated by boiling in 1X Laemmli buffer and analyzed by SDS-PAGE gel electrophoresis and was subjected to autoradiography. Alternatively, the invitro phosphorylated (by ERK2) recombinant purified NP was subjected to LC-MS/MS analysis (outsourced from V Proteomics, India).

### In vitro kinasing using analogue sensitive ERK2 followed by phosphothioester antibody mediated detection

Analogue sensitive ERK2 (ERK2 AS) was generated by introducing the Q105G mutation in the pCDNA3X-FLAG-ERK2 construct to enlarge the ATP-binding pocket, as described previously (8, 9). ERK2 or ERK2-AS were immuno-purified from transiently transfected HEK293T cells and used for in vitro kinasing as described above with only exception of using 1mM γ-thio-ATP (Abcam) or γ-thio-ATP-analogue (N6-Phenylethyl γ-thio-ATP, Biolog Life Science Institute, Bremen, Germany) instead of the mixture of γ-32P ATP. The invitro phosphorylated thio-phosphate conjugated substrates were alkylated using 2.5 mM of p-nitro benzyl mesylate (PNBM) at 22^0^C for 2 hours with intermittent shaking every 15 minutes. Finally, the reaction was stopped using 1X laemmli buffer and analysed through SDS-PAGE. The thio-phosphorylated substrates were detected through western blotting using anti-thiophosphate ester antibody [51-8] (abcam).

### In cell kinasing using analogue sensitive ERK2

HEK293T cells transiently transfected with plasmids overexpressing ERK2 or ERK2-AS or empty vector control were infected with influenza A/H1N1/WSN/1933 virus or mock infected (MOI 0.5). At 12hpi, cells were stimulated with 1.5ug/ml PMA for 30 minutes or left unstimulated. Subsequently, 1.5×10^6^ cells from each set were resuspended in ice cold 1x ERK2 kinase buffer (mentioned above) reconstituted in PBS containing complete protease and phosphatase inhibitor cocktails and 50 ug/ml digitonin (Sigma) and incubated on ice for 5 min. In cell kinasing is carried out in the same buffer (without digitonin) containing 100uM N6-phenethyl ATP-γ-S and 1 mM ATP at 30^0^C for 40 minutes with gentle rocking. Cells were then lysed in 0.5 ml RIPA buffer (50 mM Tris-HCl (pH 8), 150 mM NaCl, 1.0% NP-40 and 0.1% SDS) containing 25 mM EDTA followed by immunoprecipitation using anti-RNP antibody. The immunoprecipitated thio-phosphate conjugated NP was alkylated using 2.5 mM of p-nitro benzyl mesylate (PNBM) for 2hrs at 22^0^Cand detected through western blotting using anti-thiophosphate ester antibody [51-8] (abcam).

### Immunofluorescence assay & image analysis

A549 cells grown on coverslips were infected with WSN-PB2-FLAG at an MOI of 5 followed by fixation with 3% formaldehyde at different times of post infection as indicated for different experiments. Fixative was quenched with 0.1M Glycine and the cells were permeabilized with 0.1% Triton-X 100 in PBS for 15 min at room temperature, followed by blocking with 3% BSA in PBS for an hour in room temperature. Cells were incubated with respective primary antibodies in blocking solution for overnight at 4^0^C. NP was detected with anti-RNP primary antibody and Alexa Fluor 488-conjugated donkey anti-goat IgG secondary antibody (I:1000 dilution) (Invitrogen). ERK2/ MEK1 and PKC⍺ were detected with specific primary antibodies at recommended dilution (Cell Signaling) and Alexa Fluor 568, Alexa Flour 555-conjugated donkey anti-rabbit IgG and Anti-mouse secondary antibody (1:1000 dilution) (Invitrogen) respectively. DAPI (1μg/ml) was used to stain the nucleus. Cells were imaged using 63x and 100x oil immersion objective lenses at 405nm, 488nm and 561nm lasers in confocal microscope (Olympus FV3000) and post-processed with ImageJ software. 16-bit raw data sets were thresholded and converted to TIFF stacks for analysis.

Specific signal intensity in the nucleus and through the cell was measured through ImageJ software. 50 cells (n=50) were randomly chosen for each experimental set (at least 10 cells per field). Nucleus / total cell intensity was calculated for each cell and plotted on the Y axis of the graph. Mean intensity with standard deviation of each sets was compared by one-way ANOVA.

### Super resolution structured illumination microscopy (SIM) and image analysis

A549 cells grown on coverslips were infected with WSN-PB2-FLAG at an MOI of 5 following protocol as mentioned above in IFA. Super resolution imaging was performed on an Elyra 7 lattice SIM imaging system (ZEISS) equipped with 63x oil immersion objective, and 405nm, 488nm and 561nm diode lasers. Raw data was acquired and reconstituted as per the standard protocol of the company by using appropriate oil refractive index to generate a supper resolution Lattice SIM^2^ image (resolution excellence down to less than 60 nm) in the ZEN software (ZEISS). 32-bit reconstituted data sets were thresholded and converted to TIFF stacks for analysis.

Colocalization of NP – ERK2/MEK1/PKCα was analysed in image J (Fiji) software. Region of interest was selected with a rectangle tool and RGB profile plot was generated. Pearson correlation coefficient and Mander’s overlap coefficients M1 and M2 were calculated for the total field for each data sets.

### Immunoprecipitation assay

HEK293T cells expressing NP and other interacting partners (ERK2, MEK1 and PKCα-CAT, PKCα-CAT-DN or full length PKCα WT(PKCα-FL) were lysed in radio-immunoprecipitation assay (RIPA) buffer (50 mM Tris-HCl (pH 7.5), 150 mM NaCl, 2 mM EDTA, 1% NP-40, 0.5% deoxycholate, 0.1% SDS) supplemented with 5mg/ml of BSA and clarified by centrifugation as described (3). Lysates were incubated with appropriate antibodies and immunocomplexes were captured on Protein A sure beads (Biorad). Beads were subsequently washed once with RIPA buffer containing 500 mM NaCl and 5mg/ml BSA and finally twice in RIPA buffer without BSA. Immunoprecipitated samples were analyzed by western blotting.

A549 cells were infected with either A/WSN/1933 (H1N1) or influenza B (B/ Brisbane/ 60/ 2008) virus (MOI 10) respectively. The infected cell lysates were subsequently proceeded for immunoprecipitation assay. Lysates were incubated with anti NP antibody and subsequently captured using Easy view beads (Sigma). The beads were washed with RIPA buffer and finally the immunoprecipated complexes were analysed by western blot assay.

### Western Blot analysis

Cell lysates or immunoprecipitated samples were resolved through SDS-PAGE and transferred to Immobilon polyvinylidene difluoride (PVDF) membrane (Biorad). The membrane is blocked using 5% skim milk in TBST for 1h at room temperature and incubated with appropriate primary antibodies at 4^0^C overnight. Followed by washing, the membrane was incubated with respective secondary antibodies conjugated with horseradish peroxidase (SIGMA) and detection was carried out using SuperSignal™ West Pico PLUS Chemiluminescent Substrate (Invitrogen) in Chemidoc-MP (Biorad) multimodal gel imaging system.

### LC-MS analysis

A549 cells were infected with WSN virus (MOI 10) and at 24hpi the cells were lysed in RIPA buffer. The infected cell lysates were boiled in 5X laemmli buffer, separated by SDS-PAGE gel electrophoresis followed by Commassie blue staining. The WSN NP protein band from its desired molecular weight position was excised from the gel, cut into 3-5 gel cubes and subjected to in-gel trypsin digestion protocol. The eluted peptides were further utilized for mass spectrometric analysis. Peptides were characterized using a Thermo Q-exactive-HF-X mass spectrometer coupled to a Thermo Easy nLC 1200. Samples separated at 300nl/min on an Acclaim PEPMAP 100 trap (75uM, 2CM, c18 3um, 100A) and an easyspray 100 Column (75um, 25cm, c18, 100A) using a 120 minute gradient with an initial starting condition of 2% B buffer (0.1% formic acid in 90% Acetonitrile) and 98% A buffer (0.1% formic acid in water). Buffer B was increased to 28% over 90 minutes, then up to 40% in an additional 10 minutes. High B (90%) was run for 15 minutes afterwards. The mass spectrometer was outfitted with a Thermo nanospray easy source with the following parameters: Spray voltage: 1.8, Capillary temperature: 250dC, Funnel RF level=40. Parameters for data acquisition were as follows: for MS data the resolution was 60,000 with an AGC target of 3e6 and a max IT time of 50 ms, the range was set to 400-1600 m/z. MS/MS data was acquired with a resolution of 15,000, an AGC of 1e5, max IT of 50 ms, and the top 30 peaks were picked with an isolation window of 1.6m/z with a dynamic execution of 25s.

### Data analysis

The resulting samples were processed using Thermo Proteome Discoverer 2.20.388. The custom database was made with supplied sequences that were downloaded from UniProt and searched with the following parameters: a tryptic enzyme with a max of 2 missed cleavages, a precursor mass tolerance of 10ppm, and a fragment mass tolerance of 0.02 Da with pSTY modifications. The FDR rate was set at 0.01.

### Proximity Ligation Assay (PLA) and Immuno Fluorescence Assay (IFA)

Approximately 6×10^4^ A549 cells grown on eight well chamber glass slides were infected with influenza A virus strain, A/WSN/1933 (MOI 5). The cells were fixed with 4% Paraformaldehyde for 20 min and permeabilized with 0.2% Triton X-100 in PBS for 5 min. PLA was performed according to the manufacturer’s instructions using the following kits and reagents: Duolink In Situ PLA Probe Anti-Rabbit MINUS (Sigma-Aldrich # DUO92005), Duolink In Situ PLA Probe Anti-Goat PLUS (Sigma-Aldrich # DUO92003), Duolink In Situ Detection Reagents Red (Sigma-Aldrich # DUO92008). The cells were probed with primary ERK2 antibody raised in rabbit (Cell signalling) and RNP antibody raised in goat (BEI resources) at 37^0^C for 1h. For negative control, respective isotype matched IgG antibodies were used for the study. After performing ligation, the amplification by the polymerase was carried out for 85 min at 37^0^C. Thereafter, the cells were washed twice with Wash Buffer B for 10 min and then the samples were processed for IFA using the protocol as mentioned before using anti NP antibody (H16/L10) raised in mouse (BEI resources). PLA signals were detected as distinct fluorescent dots or puncta at 640nm wavelength using Olympus FV3000 confocal microscope at 63x oil immersion objective lense. 16-bit raw data sets were thresholded and converted to TIFF stacks for analysis using imageJ analysis software provided by the respective manufacturers. Number of PLA dots were counted for 100 cells by ImageJ software for each experimental set and plotted on the Y-axis of the graph. Average number of dots with standard deviation of each set was compared by one-way ANOVA.

Intensity of the PLA and NP signal, in the nucleus and throughout the cell were measured by ImageJ. Intensity ratio (Nucleus / Nucleus +Cytoplasm) was plotted to show their colocalization. Mean intensity with standard deviation of each sets were compared by two-way ANOVA.

### Protein expression, purification and oligomerization state determination

WT or mutant NP genes cloned in pET28a vector were expressed in *E*. *coli* strain BL21 (DE3) (Novagen) and purified using Ni-NTA affinity chromatography (Qiagen). Purified proteins were treated with RNaseA and further purified through a HiTrap *Heparin* HP column (GE Healthcare). Proteins were concentrated to 4 mg/ml and incubated at 4°C for 96 hours in buffer containing 50mM Tris, pH7.5, 200mM NaCl and 1mM TCEP. Subsequently, the oligomerization state of the proteins were analysed through size-exclusion chromatography using ENrich™ SEC 650 10 x 300 Column (Biorad) calibrated with appropriate molecular weight standards.

### RNA extraction and qRT-PCR

Total RNA was isolated from appropriately treated cells using the Trizol reagent (Invitrogen) following manufacturer’s instructions. 3μg RNA was reverse transcribed by using Moloney murine leukemia virus (M-MuLV) reverse transcriptase enzyme according to the manufacturer’s instructions. For qPCR, the synthesized 2ul of cDNA was used as a template with iTaq Universal SYBR Green Supermix (Bio-Rad). The cDNAs used for qPCR analysis was prepared using Power SYBR Green PCR Master Mix (Bio-Rad, # 172-5125) in a QuantStudio5 Applied Biosystems real-time PCR using the Influenza A/H1N1 NP segment specific primers either targeting the negative sense genomic RNA or mRNA. All RNA levels were normalized to GAPDH mRNA levels and calculated as the delta-delta threshold cycle (ΔΔCT).

### Polymerase activity assay

HEK293T cells were transfected with pCMV-PA, pCMV-PB1, pCDNA-PB2-3xFLAG, pcDNA-NP-V5 or mutant pcDNA-NP-V5 plasmids and pHH21vNA-Luc reporter plasmids by using Lipofectamine-3000. Cells were lysed 24 hours post-transfection in cell culture lysis reagent (CCLR) (Promega), and luciferase activity was measured using the luciferase assay system luminometer-Glomax 20/20 (Promega). Expression of WSN RNP components were analyzed by Western blotting.

### Statistical analysis

All experiments were performed in triplicates, and each data was repeated at least three times except for the epitope mapping experiments, which were performed in duplicates. Graphs are plotted using Graphpad Prism 10.1.2 and represented as mean standard deviations (n = 3). Results were compared by performing a two-tailed Student’s t-test and one-way ANOVA. Significance is defined as P < 0.05, and statistical significance is indicated with an asterisk (*). *P < 0.001 were considered statistically significant. Significance was denoted as, (ns > 0.05; *P `≤` 0.05; **P `≤` 0.01; ***P `≤` 0.001) and considered to be statistically significant.

## Supplementary Figures

**Fig S1.**
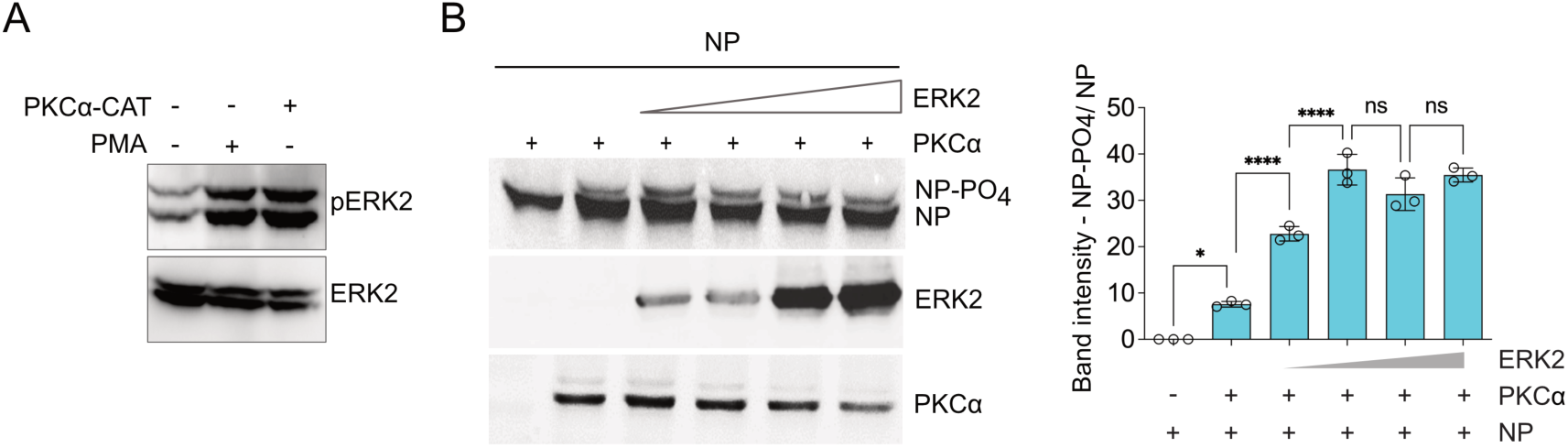
(A) HEK293T cells either mock treated or treated with PMA or transfected with PKCα-CAT were analyzed for ERK2 activation by western blot analysis using ERK2 or phospho-ERK2 antibody. (B) Cells overexpressing NP-V5 with PKCα-CAT along with increasing concentration of ERK2-FLAG were analyzed by western blot analysis using anti-V5 antibody.

**Fig S2.**
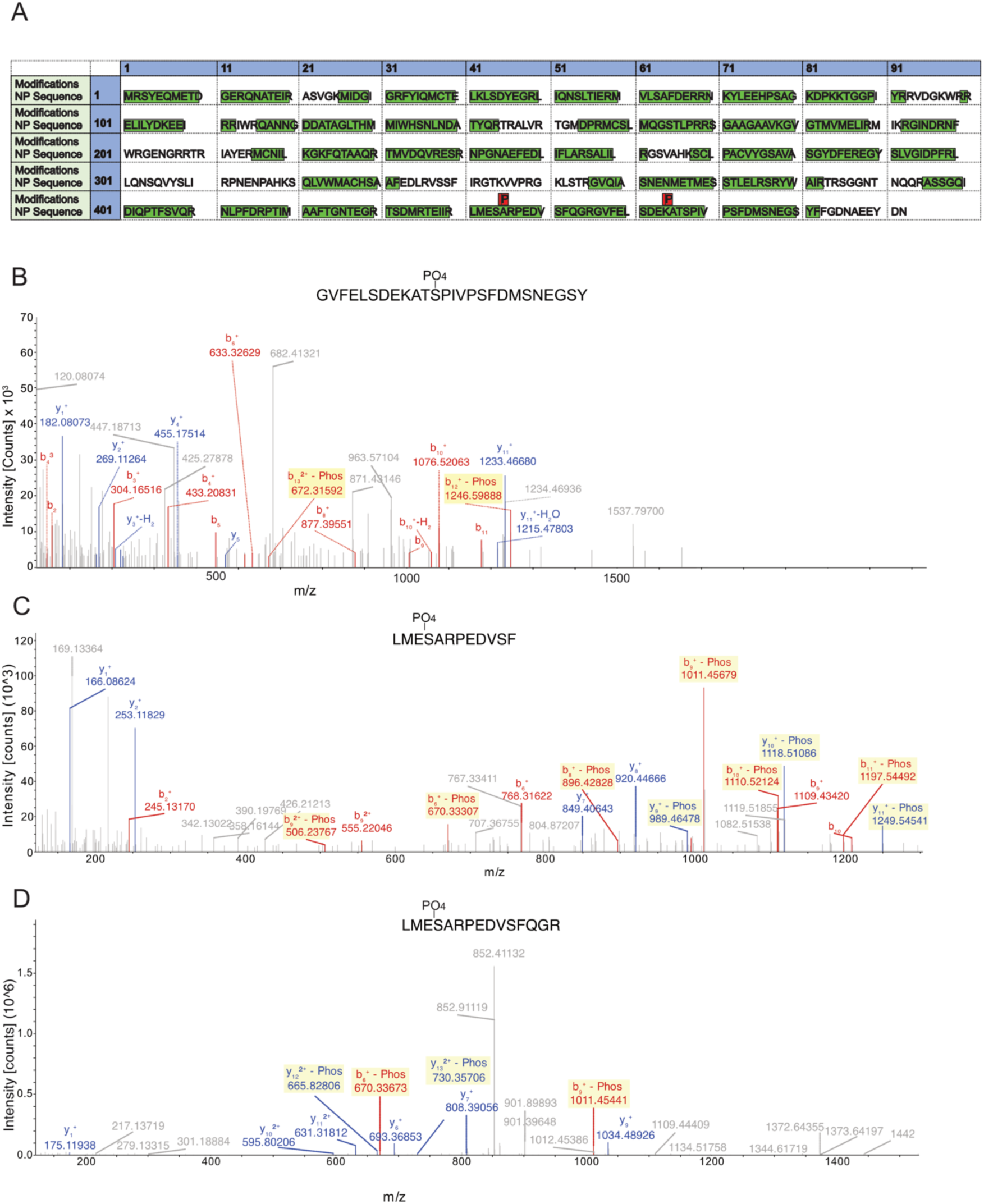
In vitro phosphorylated NP was subjected to LC-MS/MS analysis. (A) Table showing peptide coverage for NP with phosphorylation of specific amino acid residues. (B-D) showing chromatograms of the phsopho-peptides subjected to CID fragmentation. Y and B ions are labelled in blue and red respectively.

**Fig S3.**
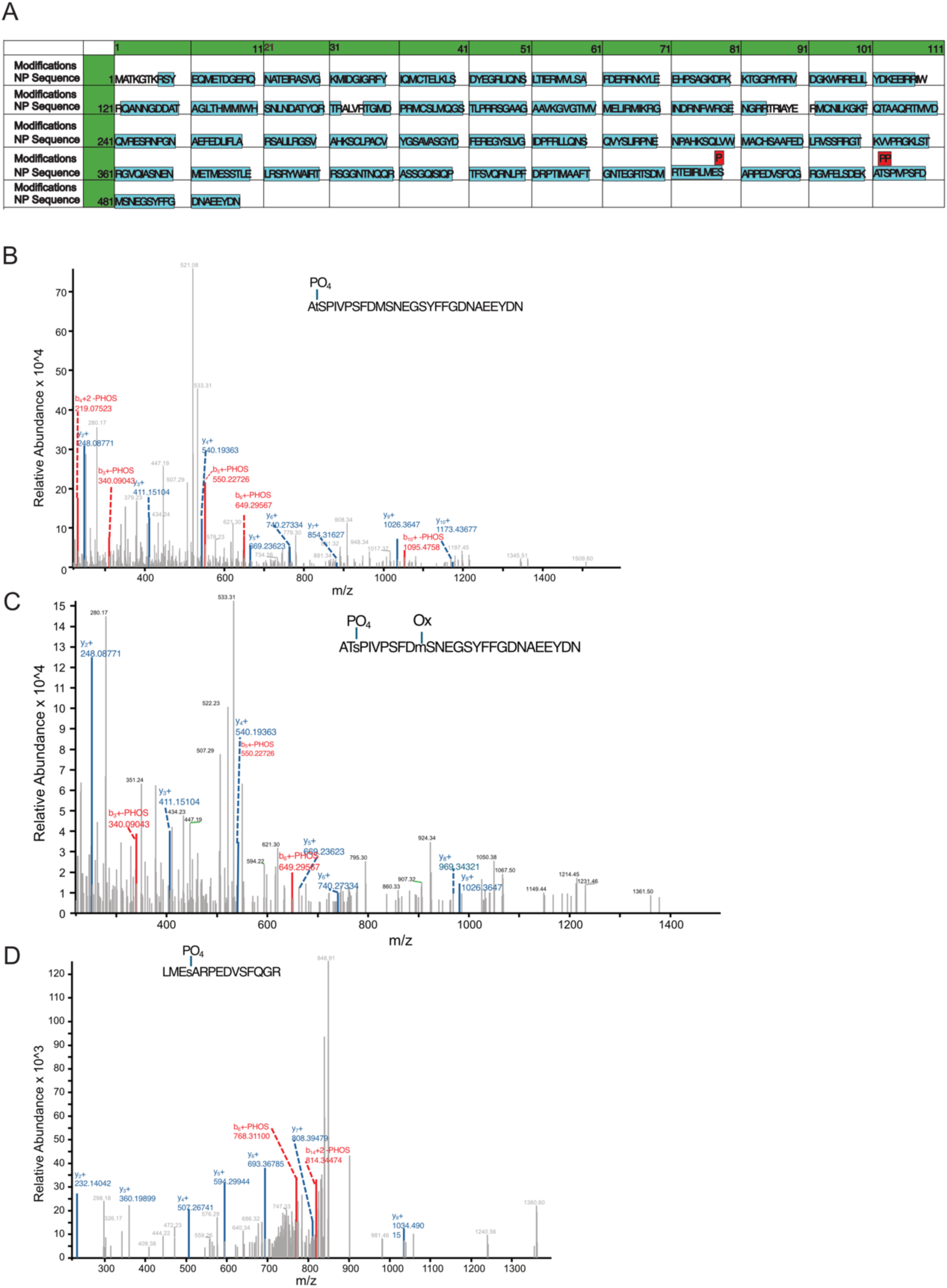
NP purified from A/WSN/1933 virus infected cells was subjected to LC-MS/MS analysis. (A) Table showing peptide coverage for NP with phosphorylation of specific amino acid residues. (B-D) showing chromatograms of the phsopho-peptides subjected to CID fragmentation. Y and B ions are labelled in blue and red respectively.

**Fig S4.**
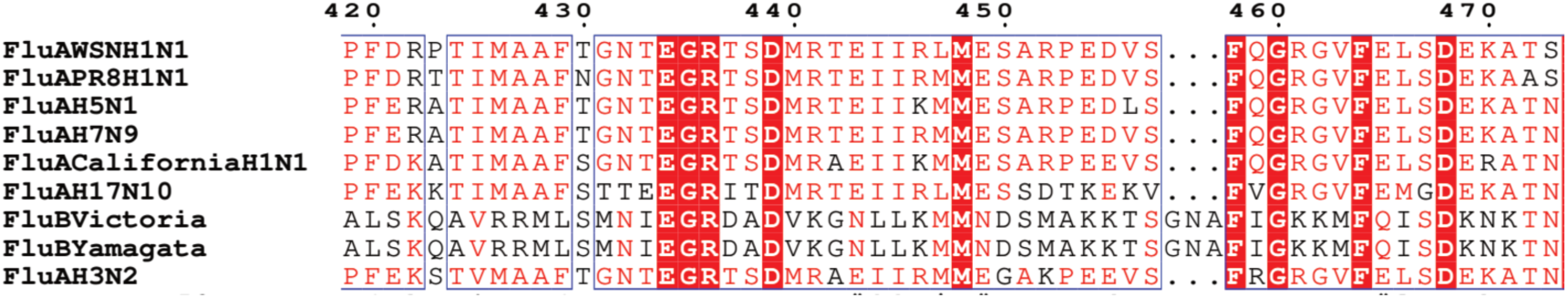
Multiple sequence alignment of NP of different influenza A and B viruses.

**Fig S5.**
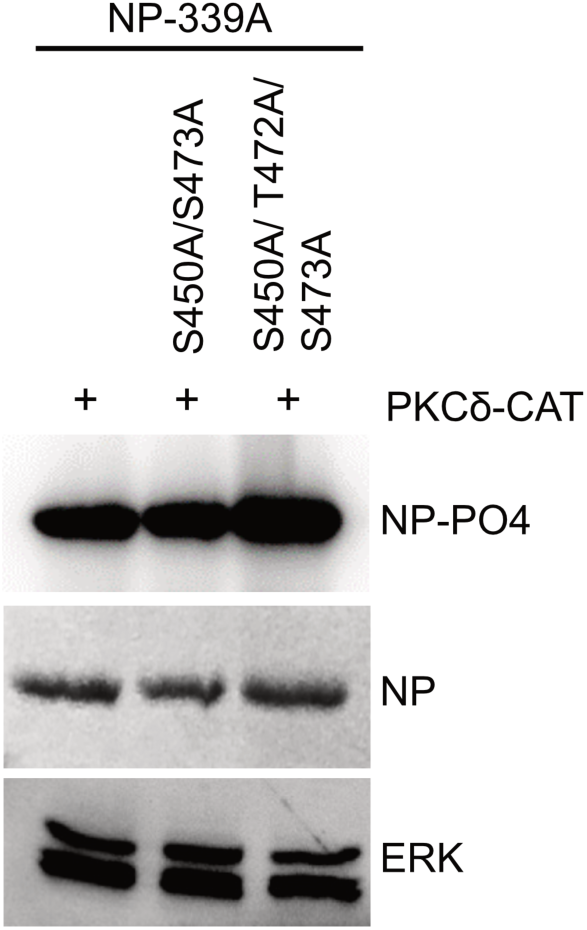
Monomeric NP-E339A harboring various phosphonull alanine mutations were subjected to in vitro kinasing by -PKCδ-CAT.

**Fig S6.**
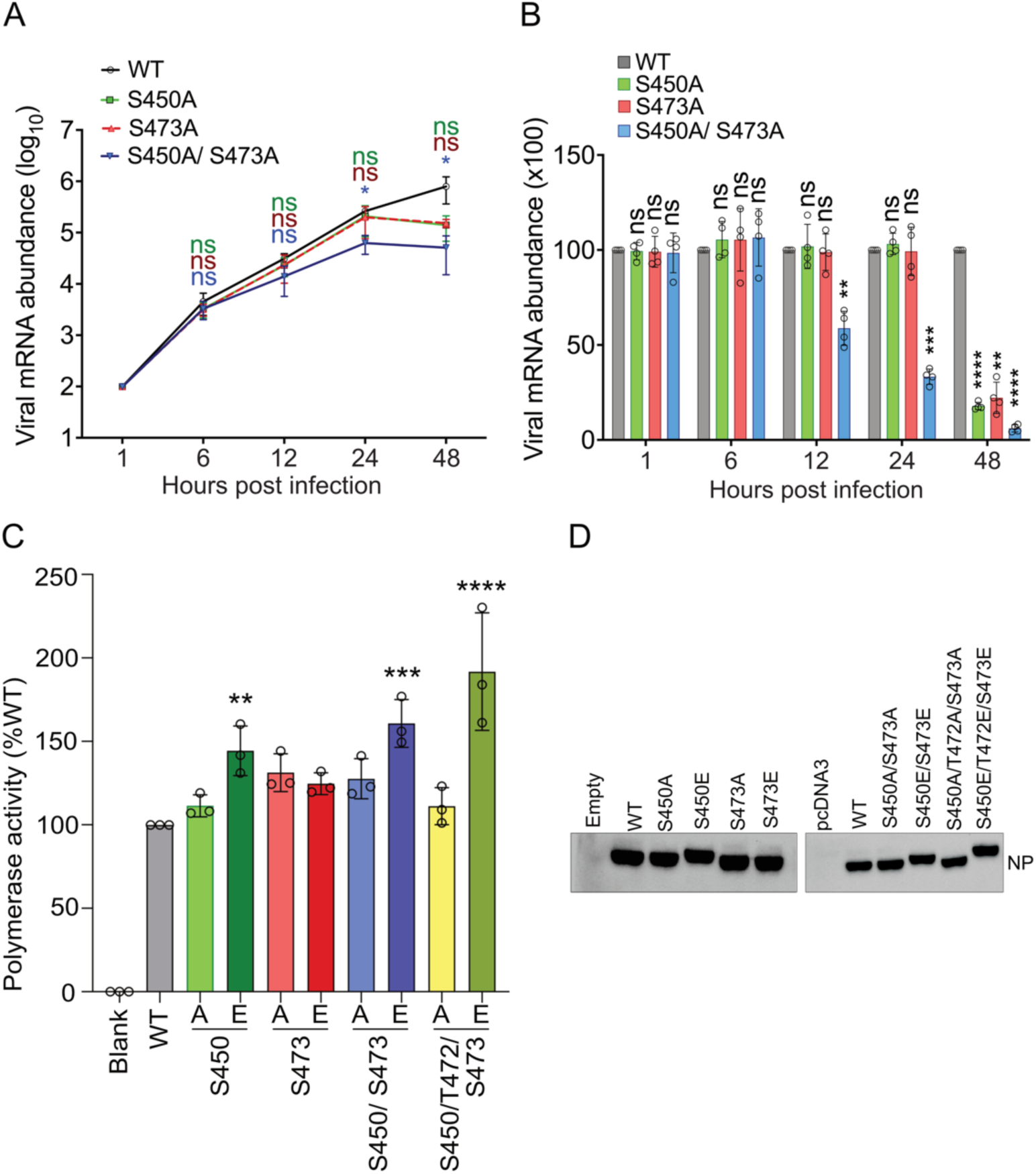
A549 cells were infected with WT and mutant viruses harboring phospho-null alanine substitutions at the ERK2 phosphorylation sites. Absolute (A) and relative (B) abundance of viral mRNA at different times of post infection were quantified using realtime qRT-PCR. (C) Ability of the WT and mutant NP proteins to support RNP activity and viral RNA synthesis using reporter RNPs with positive sense viral RNA (cRNA) template reconstituted in HEK293T cells through transient transfection. (D) Expression levels of the WT and mutant NP proteins assessed through western blot analysis.

**Fig S7.**
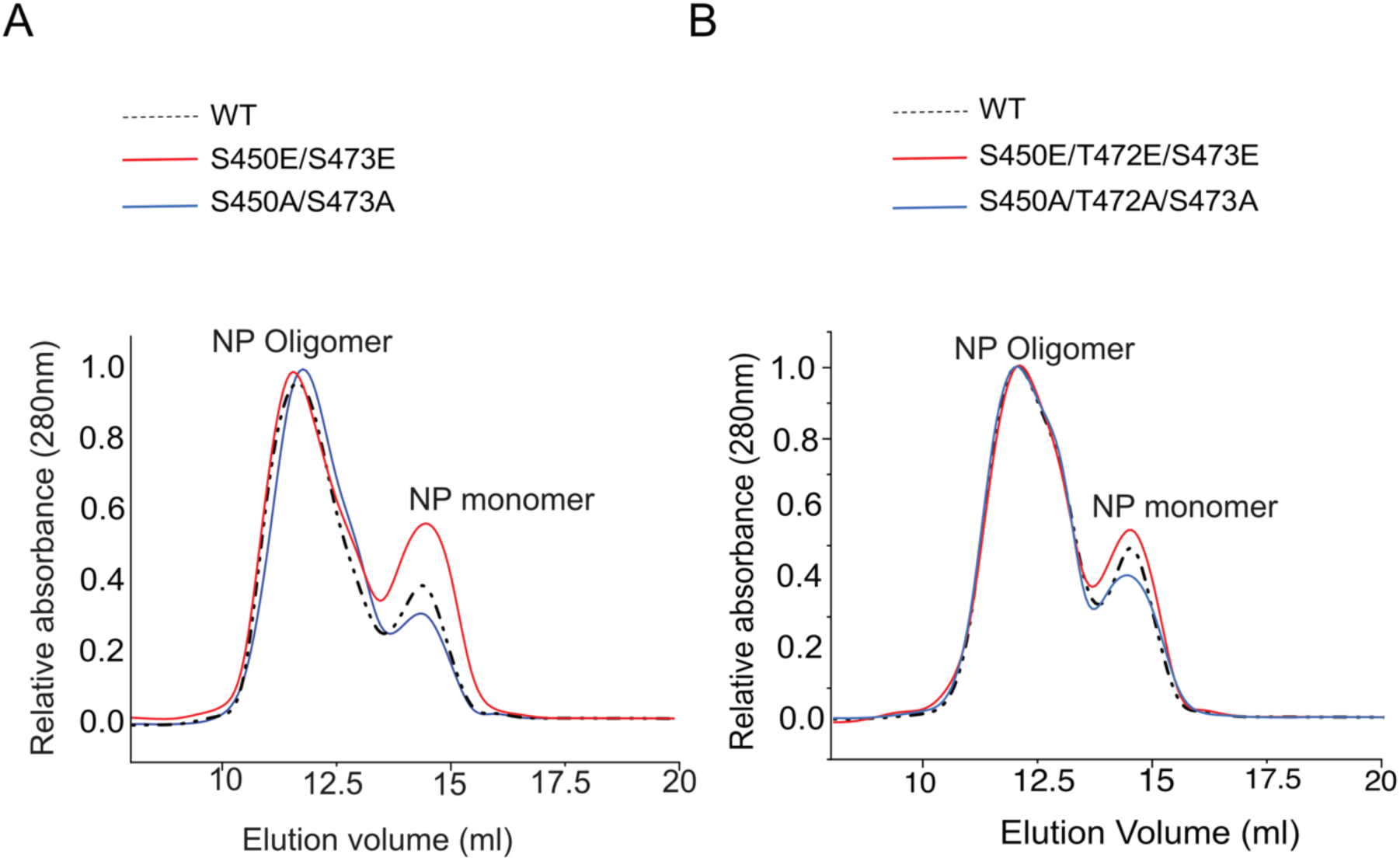
WT and mutant NP proteins, expressed and purified from bacteria were subjected to extensive RNaseA treatment followed by size exclusion analysis to evaluate their oligomerization potential. (A) showing oligomerization profile of the NP S450A/S473A and NP S450E/S473E and (B) showing S450A/ T472A/ S473A and NP S450E/ T472E/ S473E mutant proteins in comparison to the WT.

**Fig S8.**
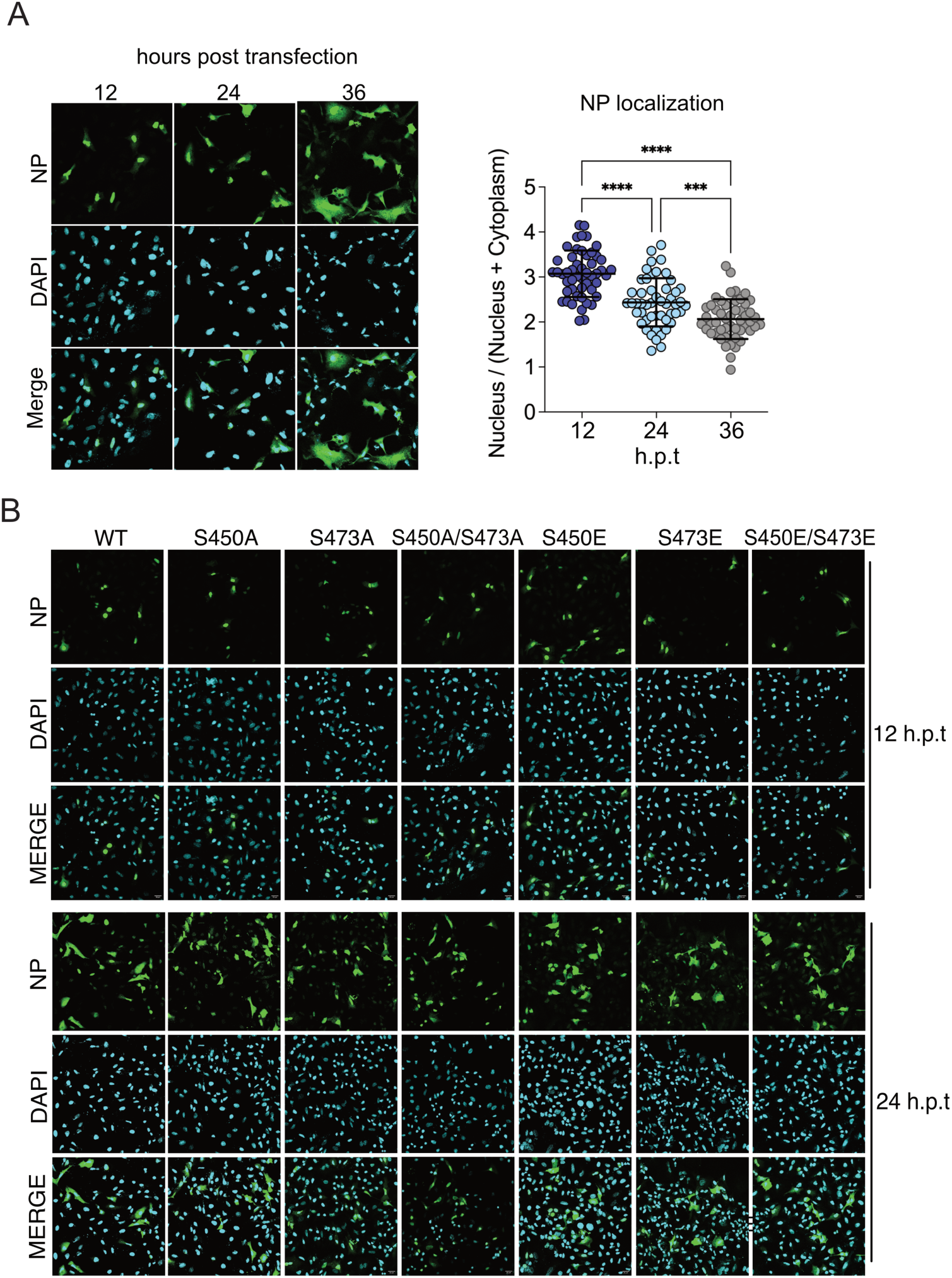
(A) Influenza A/H1N1 NP was expressed in A549 cells through transient transfection, fixed and stained using specific antibody at 12, 24, 36 hours post transfection. Imaged using confocal microscope. 50 cells from 5 different filed was analysed using image J software to present nucelar-cytoplasmic distributions of NP. (B) WT and mutant NP proteins expressed in A549 cells were fixed and immuno-stained with anti-NP antibody at 12 and 24 hpt. Cells were imaged using confocal microscope.

**Fig S9.**
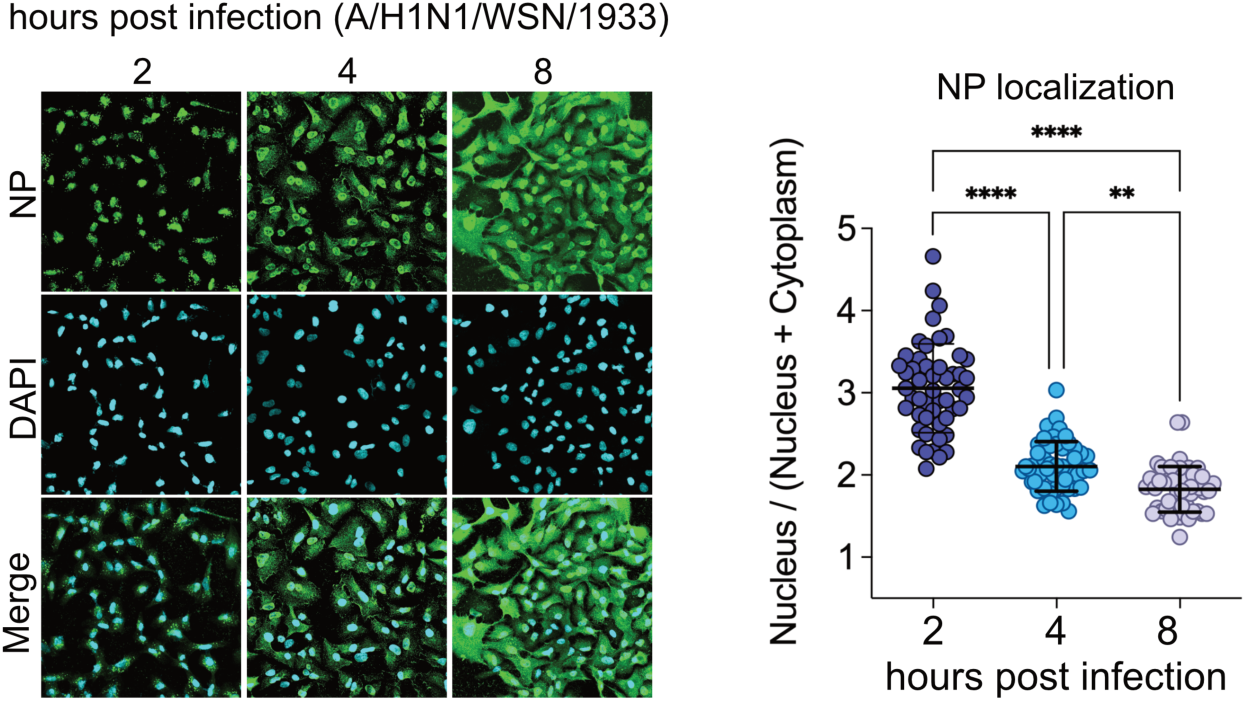
A549 cells infected with WT A/WSN/1933 viruses (MOI:5). 2, 4 and 8 hpicells were fixed and immune-stained using anti-RNP antibody and imaged using confocal microscope. Nucleo-cytoplasmic distribution of NP was analysed using imageJ software.

**Fig S10.**
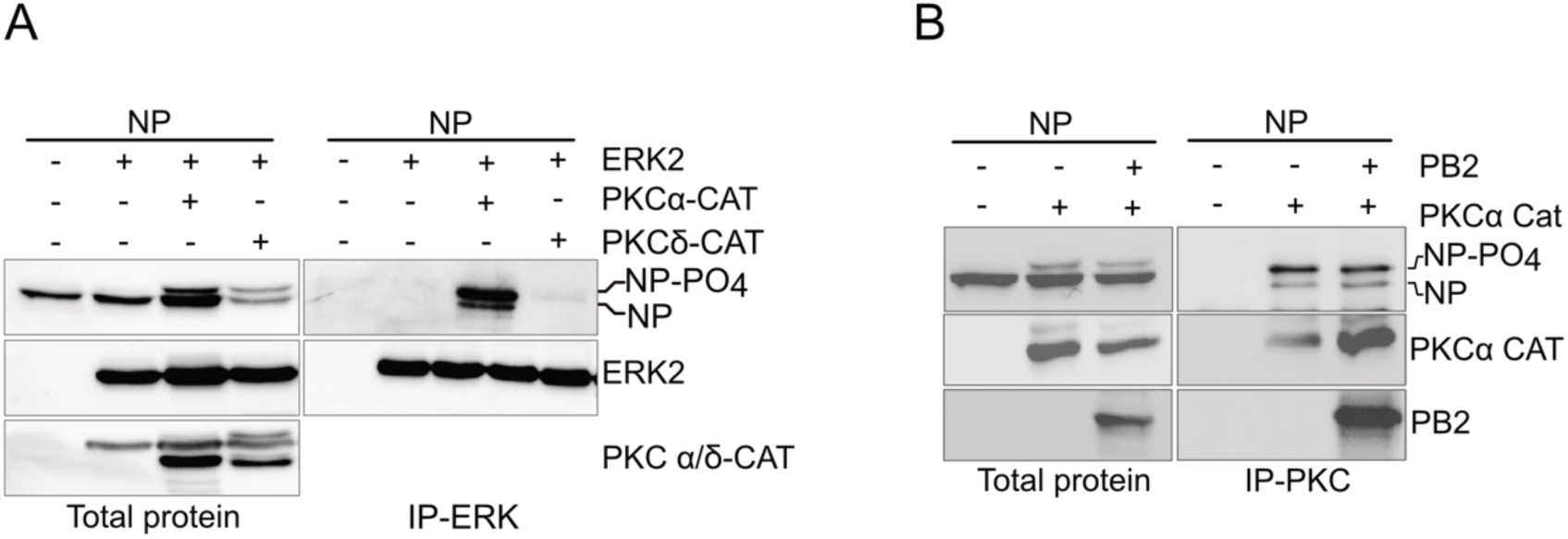
(A) Cells were transiently transfected to coexpress NP and ERK2 either in the context of PKC-α-CAT or PKC-δ-CAT overexpression. Immunoprecipitated using ERK2 specific antibody and coprecipitation was observed with NP specific antibody. (B) Co-IP of NP with PKC-α-CAT either in the presence or absence of PB2.

**Fig S11.**
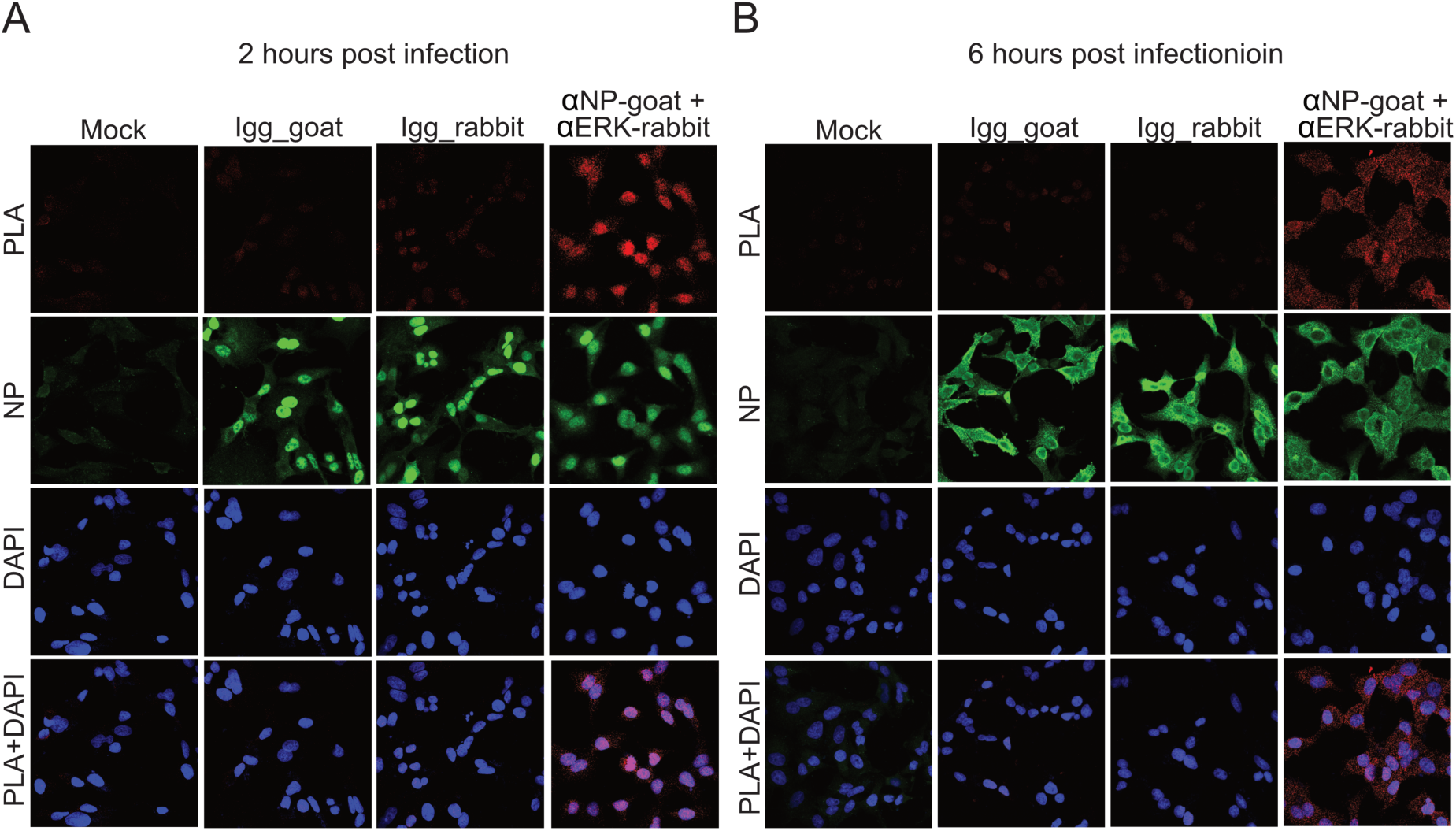
Cells infected with influenza A/H1N1 virus or mock infected were subjected to PLA using anti-RNP (goat) and anti-ERK2 (rabbit) primary antibodies or with one of the primary antibody along with the IgG isotype control for the other. Post PLA, cells are stained with anti-NP antibody (mouse) and imaged using confocal microscope.

---

**Table S1.** Ion series for the peptide - LMESARPEDVSF.

| #1 | $b^+$ | $b^{2+}$ | Seq. | $y^+$ | $y^{2+}$ | #2 |
| --- | --- | --- | --- | --- | --- | --- |
| 1 | 114.09134 | 57.54931 | L |  |  | 12 |
| 2 | 245.13183 | 123.06955 | M | 1347.52866 | 674.26797 | 11 |
| 3 | 374.17442 | 187.59085 | E | 1216.48817 | 608.74773 | 10 |
| 4 | 541.17278 | 271.09003 | S-Phospho | 1087.44558 | 544.22643 | 9 |
| 5 | 612.20989 | 306.60858 | A | 920.44722 | 460.72725 | 8 |
| 6 | 768.31100 | 384.65914 | R | 849.41011 | 425.20869 | 7 |
| 7 | 865.36377 | 433.18552 | P | 693.30900 | 347.15814 | 6 |
| 8 | 994.40636 | 497.70682 | E | 596.25623 | 298.63175 | 5 |
| 9 | 1109.43330 | 555.22029 | D | 467.21364 | 234.11046 | 4 |
| 10 | 1208.50172 | 604.75450 | V | 352.18670 | 176.59699 | 3 |
| 11 | 1295.53374 | 648.27051 | S | 253.11828 | 127.06278 | 2 |
| 12 |  |  | F | 166.08626 | 83.54677 | 1 |

**Table S2.** Ion series for the peptide: LMESARPEDVSFQGR.

| #1 | $b^+$ | $b^{2+}$ | Seq. | $y^+$ | $y^{2+}$ | #2 |
| --- | --- | --- | --- | --- | --- | --- |
| 1 | 114.09134 | 57.54931 | L |  |  | 15 |
| 2 | 245.13183 | 123.06955 | M | 1688.70981 | 844.85854 | 14 |
| 3 | 374.17442 | 187.59085 | E | 1557.66933 | 779.33830 | 13 |
| 4 | 541.17278 | 271.09003 | S-Phospho | 1428.62673 | 714.81700 | 12 |
| 5 | 612.20989 | 306.60858 | A | 1261.62837 | 631.31783 | 11 |
| 6 | 768.31100 | 384.65914 | R | 1190.59126 | 595.79927 | 10 |
| 7 | 865.36377 | 433.18552 | P | 1034.49015 | 517.74871 | 9 |
| 8 | 994.40636 | 497.70682 | E | 937.43739 | 469.22233 | 8 |
| 9 | 1109.43330 | 555.22029 | D | 808.39479 | 404.70103 | 7 |
| 10 | 1208.50172 | 604.75450 | V | 693.36785 | 347.18756 | 6 |
| 11 | 1295.53374 | 648.27051 | S | 594.29944 | 297.65336 | 5 |
| 12 | 1442.60216 | 721.80472 | F | 507.26741 | 254.13734 | 4 |
| 13 | 1570.66074 | 785.83401 | Q | 360.19899 | 180.60314 | 3 |
| 14 | 1627.68220 | 814.344744 | G | 232.14042 | 116.57385 | 2 |
| 15 |  |  | R | 175.11895 | 88.06311 | 1 |

**Table S3.** Ion series for the peptide: GVFELSDEKATSPIVPSFDMSNEGSY.

| #1 | $b^+$ | $b^{2+}$ | $b^{3+}$ | Seq. | $y^+$ | $y^{2+}$ | $y^{3+}$ | #2 |
| --- | --- | --- | --- | --- | --- | --- | --- | --- |
| 1 | 58.02874 | 29.51801 | 20.01443 | G |  |  |  | 26 |
| 2 | 157.09715 | 79.05222 | 53.03724 | V | 2829.22125 | 1415.11426 | 943.74527 | 25 |
| 3 | 304.16557 | 152.58642 | 102.06004 | F | 2730.15284 | 1365.58006 | 910.72246 | 24 |
| 4 | 433.20816 | 217.10772 | 145.07424 | E | 2583.08442 | 1292.04585 | 861.69966 | 23 |
| 5 | 546.29223 | 273.64975 | 182.76893 | L | 2454.04183 | 1227.52455 | 818.68546 | 22 |
| 6 | 633.32425 | 317.16576 | 211.77960 | S | 2340.95776 | 1170.98252 | 780.99077 | 21 |
| 7 | 748.35120 | 374.67924 | 250.12192 | D | 2253.92574 | 1127.46651 | 751.98010 | 20 |
| 8 | 877.39379 | 439.20053 | 293.13611 | E | 2138.89879 | 1069.95303 | 713.63778 | 19 |
| 9 | 1005.48875 | 503.24801 | 335.83444 | K | 2009.85620 | 1005.43174 | 670.62358 | 18 |
| 10 | 1076.52587 | 538.76657 | 359.51347 | A | 1881.76124 | 941.38426 | 627.92526 | 17 |
| 11 | 1177.57354 | 589.29041 | 393.19603 | T | 1810.72412 | 905.86570 | 604.24623 | 16 |
| 12 | 1344.57190 | 672.78959 | 448.86215 | S-phospho | 1709.67645 | 855.34186 | 570.56367 | 15 |
| 13 | 1441.62467 | 721.31597 | 481.21307 | P | 1542.67809 | 771.84268 | 514.89755 | 14 |
| 14 | 1554.70873 | 777.85800 | 518.90776 | I | 1445.62532 | 723.31630 | 482.54663 | 13 |
| 15 | 1653.77715 | 827.39221 | 551.93057 | V | 1332.54126 | 666.77427 | 444.85194 | 12 |
| 16 | 1750.82991 | 875.91859 | 584.28149 | P | 1233.47284 | 617.24006 | 411.82913 | 11 |
| 17 | 1837.86194 | 919.43461 | 613.29216 | S | 1136.42008 | 568.71368 | 379.47821 | 10 |
| 18 | 1984.93035 | 992.96881 | 662.31497 | F | 1049.38805 | 525.19766 | 350.46754 | 9 |
| 19 | 2099.95729 | 1050.48229 | 700.65728 | D | 902.31964 | 451.66346 | 301.44473 | 8 |
| 20 | 2230.99778 | 1116.00253 | 744.33744 | M | 787.29270 | 394.14999 | 263.10242 | 7 |
| 21 | 2318.02981 | 1159.51854 | 773.34812 | S | 656.25221 | 328.62974 | 219.42225 | 6 |
| 22 | 2432.0723 | 1216.54001 | 811.36243 | N | 569.22018 | 285.11373 | 190.41158 | 5 |
| 23 | 2561.11533 | 1281.06130 | 854.37663 | E | 455.17725 | 228.09227 | 152.39727 | 4 |
| 24 | 2618.13679 | 1309.57203 | 873.38378 | G | 326.13466 | 163.57097 | 109.38307 | 3 |
| 25 | 2705.16882 | 1353.08805 | 902.39446 | S | 269.11320 | 135.06024 | 90.37592 | 2 |
| 26 |  |  |  | Y | 182.08117 | 91.54422 | 61.36524 | 1 |

**Table S4:** Ion series for the peptide: ATSPIVPSFDMSNEGSYFFGDNAEEYDN, T2-Phospho (79.96633Da) Charge: +3, Monoisotopic m/z: 1061.75223 Da.

| #1 | b<sup>+</sup> | b<sup>2+</sup> | Seq. | y<sup>+</sup> | y<sup>2+</sup> | #2 |
| --- | --- | --- | --- | --- | --- | --- |
| 1 | 72.04439 | 36.52583 | A |  |  | 28 |
| 2 | 253.0584 | 127.03284 | T-Phospho | 3112.2078 | 1556.6075 | 27 |
| 3 | 340.09043 | 170.54885 | S | 2931.1938 | 1466.1005 | 26 |
| 4 | 437.14319 | 219.07523 | P | 2844.1618 | 1422.5845 | 25 |
| 5 | 550.22726 | 275.61727 | I | 2747.109 | 1374.0581 | 24 |
| 6 | 649.29567 | 325.15147 | V | 2634.0249 | 1317.5161 | 23 |
| 7 | 746.34843 | 373.67786 | P | 2534.9565 | 1267.9819 | 22 |
| 8 | 833.38046 | 417.19387 | S | 2437.9037 | 1219.4555 | 21 |
| 9 | 980.44888 | 490.72808 | F | 2350.8717 | 1175.9395 | 20 |
| 10 | 1095.4758 | 548.24155 | D | 2203.8033 | 1102.4053 | 19 |
| 11 | 1226.5163 | 613.76179 | M | 2088.7764 | 1044.8918 | 18 |
| 12 | 1313.5483 | 657.2778 | S | 1957.7359 | 979.37157 | 17 |
| 13 | 1427.5913 | 714.29927 | N | 1870.7038 | 935.85556 | 16 |
| 14 | 1556.6339 | 778.82056 | E | 1756.6609 | 878.8341 | 15 |
| 15 | 1613.6553 | 807.3313 | G | 1627.6183 | 814.3128 | 14 |
| 16 | 1700.6873 | 850.84731 | S | 1570.5969 | 785.80207 | 13 |
| 17 | 1863.7507 | 932.37897 | Y | 1483.5648 | 742.28605 | 12 |
| 18 | 2010.8191 | 1005.9132 | F | 1320.5015 | 660.75439 | 11 |
| 19 | 2157.8875 | 1079.4474 | F | 1173.4331 | 587.22018 | 10 |
| 20 | 2214.909 | 1107.9581 | G | 1026.3647 | 513.68598 | 9 |
| 21 | 2329.9359 | 1165.4716 | D | 969.34321 | 485.17524 | 8 |
| 22 | 2443.9788 | 1222.4931 | N | 854.31627 | 427.66177 | 7 |
| 23 | 2515.016 | 1258.0116 | A | 740.27334 | 370.64031 | 6 |
| 24 | 2644.0585 | 1322.5329 | E | 669.23623 | 335.12175 | 5 |
| 25 | 2773.1011 | 1387.0542 | E | 540.19363 | 270.60045 | 4 |
| 26 | 2936.1645 | 1468.5859 | Y | 411.15104 | 206.07916 | 3 |
| 27 | 3051.1914 | 1526.0993 | D | 248.08771 | 124.54749 | 2 |
| 28 |  |  | N | 133.06077 | 67.03402 | 1 |

**Table S5:** Ion series for the peptide: ATSPIVPSFDMSNEGSYFFGDNAEEYDN, S3-Phospho (79.96633 Da)

| #1 | b<sup>+</sup> | b<sup>+</sup>-P | Seq. | y<sup>+</sup> | #2 |
| --- | --- | --- | --- | --- | --- |
| 1 | 72.04439 |  | A |  | 28 |
| 2 | 173.09207 |  | T | 3128.2027 | 27 |
| 3 | 340.09043 | 242.11353 | S-Phospho | 3027.15502 | 26 |
| 4 | 437.14319 | 339.1663 | P | 2860.15666 | 25 |
| 5 | 550.22726 | 452.25036 | I | 2763.1039 | 24 |
| 6 | 649.29567 | 551.31877 | V | 2650.01983 | 23 |
| 7 | 746.34843 | 648.37154 | P | 2550.95142 | 22 |
| 8 | 833.38046 | 735.40357 | S | 2453.89866 | 21 |
| 9 | 980.44888 | 882.47198 | F | 2366.86663 | 20 |
| 10 | 1095.47582 | 997.49892 | D | 2219.79821 | 19 |
| 11 | 1242.51122 | 1144.53432 | M-Oxidation | 2104.77127 | 18 |
| 12 | 1329.54325 | 1231.56635 | S | 1957.73587 | 17 |
| 13 | 1443.58617 | 1345.60928 | N | 1870.70384 | 16 |
| 14 | 1572.62877 | 1474.65187 | E | 1756.66092 | 15 |
| 15 | 1629.65023 | 1531.67334 | G | 1627.61832 | 14 |
| 16 | 1716.68226 | 1618.70536 | S | 1570.59686 | 13 |
| 17 | 1879.74559 | 1781.76869 | Y | 1483.56483 | 12 |
| 18 | 2026.814 | 1928.83711 | F | 1320.5015 | 11 |
| 19 | 2173.88242 | 2075.90552 | F | 1173.43309 | 10 |
| 20 | 2230.90388 | 2132.92698 | G | 1026.36467 | 9 |
| 21 | 2345.93082 | 2247.95393 | D | 969.34321 | 8 |
| 22 | 2459.97375 | 2361.99685 | N | 854.31627 | 7 |
| 23 | 2531.01086 | 2433.03397 | A | 740.27334 | 6 |
| 24 | 2660.05346 | 2562.07656 | E | 669.23623 | 5 |
| 25 | 2789.09605 | 2691.11915 | E | 540.19363 | 4 |
| 26 | 2952.15938 | 2854.18248 | Y | 411.15104 | 3 |
| 27 | 3067.18632 | 2969.20943 | D | 248.08771 | 2 |
| 28 |  |  | N | 133.06077 | 1 |

**Table S6:**
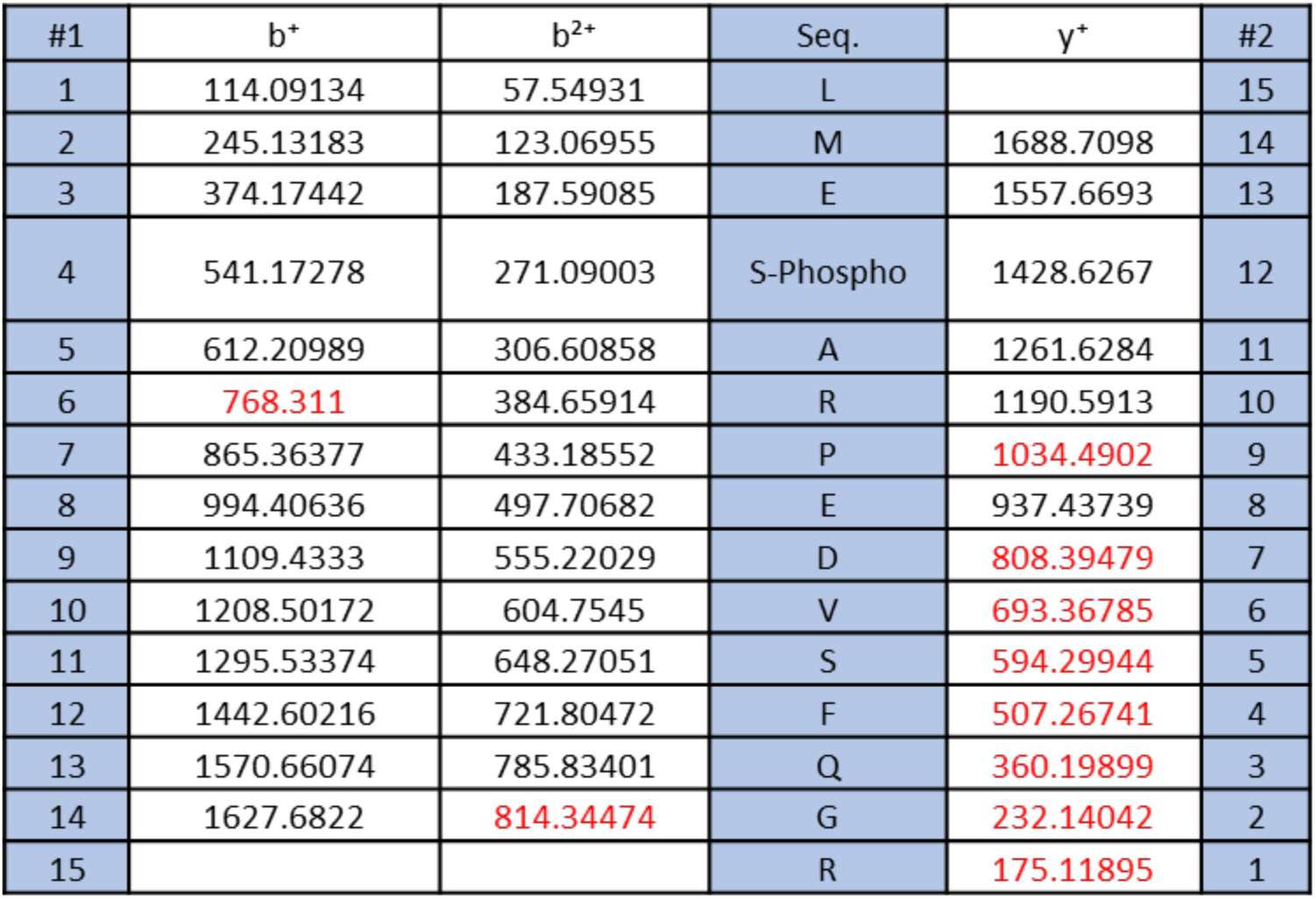
Ion series for the peptide: LMESARPEDVSFQGR, S4-Phospho (79.96633 Da)

**Table S7:** Percentage conservation of the ERK2 phosphorylation sites in different influenza virus NP proteins.

| Residue No.<br>(As per A/H1N1/WSN/1933) | Subtypes | % Conservation |
| --- | --- | --- |
| S450 | Influenza A-H1N1 | 97.01652 |
|  | Influenza A-H5N1 | 100 |
|  | Influenza A-H7N9 | 100 |
|  | Influenza A-H3N2 | 9.725251 |
| S507 | Influenza B | 99.94 |
| S473 | A-H1N1 | 1.70394 |
|  | A-H5N1 | 0 |
|  | A-H7N9 | 0 |
|  | A-H3N2 | 0.65388 |
| T472 | A-H1N1 | 95 |
|  | A-H5N1 | 100 |
|  | A-H7N9 | 100 |
|  | A-H3N2 | 46.7 |
| T531 | Influenza B | 97.21 |

**Table S8:**
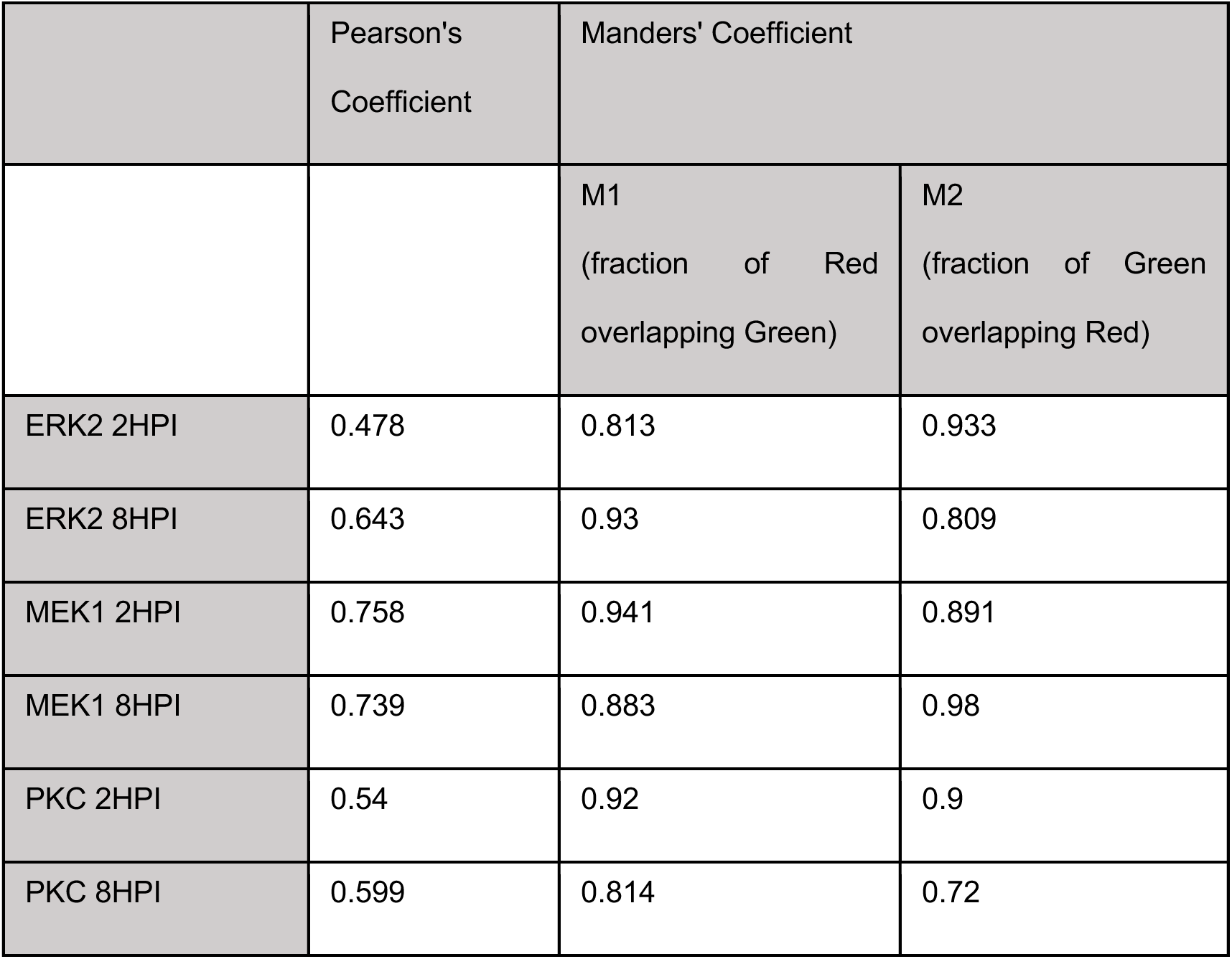
Coefficient of Co-localization from super resolution microscopy of PKC, MEK1 and ERK2 (in red) with NP (in green).

|  | Pearson's Coefficient | Manders' Coefficient |  |
| --- | --- | --- | --- |
|  |  | M1<br>(fraction of Red<br>overlapping Green) | M2<br>(fraction of Green<br>overlapping Red) |
| ERK2 2HPI | 0.478 | 0.813 | 0.933 |
| ERK2 8HPI | 0.643 | 0.93 | 0.809 |
| MEK1 2HPI | 0.758 | 0.941 | 0.891 |
| MEK1 8HPI | 0.739 | 0.883 | 0.98 |
| PKC 2HPI | 0.54 | 0.92 | 0.9 |
| PKC 8HPI | 0.599 | 0.814 | 0.72 |

